# VRC01 Selects Rare HIV Escape Mutations After Acquisition in Antibody-Mediated Prevention Trials

**DOI:** 10.1101/2025.10.29.685411

**Authors:** Carolyn Williamson, Lyle Curry, Nonhlanhla N. Mkhize, Elena E. Giorgi, Craig A. Magaret, Bronwen E. Lambson, Sinethemba Bhebhe, Haajira Kaldine, Thandeka Moyo-Gwete, Morgane Rolland, Raabya Rossenkhan, Nina Marie G. Garcia, Chivonne Moodley, Anna Yssel, Yunda Huang, Alaine A. Marsden, Daniel B. Reeves, Bryan T. Mayer, Roger E. Bumgarner, Nicolas Beaume, Dylan H. Westfall, Michal Juraska, Allan C. DeCamp, Hugh Murrell, Hongjun Bai, Wenjie Deng, Alec P. Pankow, Tanmoy Bhattacharya, Talita York, Nonkululeko Ndabambi, Lennie Chen, Hong Zhao, Asanda Gwashu-Nyangiri, Ruwayhida Thebus, Paula Cohen, Ben Murrell, Shelly Karuna, John Hural, Nyaradzo Mgodi, Srilatha Edupuganti, Lynn Morris, David Montefiori, M. Juliana McElrath, Myron S. Cohen, Lawrence Corey, Paul T. Edlefsen, Peter B. Gilbert, Penny L. Moore, James I. Mullins

**Author notes:** DHW and WD: Vaccine and Infectious Disease Division, Fred Hutchinson Cancer Center, Seattle, WA, USA. Contributed equally to this article (Lyle Curry, Nonhlanhla N. Mkhize, Elena E. Giorgi). Contributed equally to this article (Penny L. Moore, James I. Mullins).

## Abstract

Broadly neutralizing antibodies (bnAbs) show promise in HIV prevention, yet viral escape remains a challenge. In the Antibody Mediated Prevention (AMP) trials, the CD4 binding site (CD4bs) bNAb VRC01 blocked acquisition by VRC01-sensitive strains. However, its influence on viral evolution post-acquisition is not fully understood. Here we analyzed >12,000 HIV env sequences from 47 participants from the AMP trials, identifying VRC01-mediated *de novo* escape mutations in 8 of 26 VRC01-treated participants but none in 21 placebo participants. These mutations were found at very low frequency (<1%) in global viruses. Escape mutations, primarily located in the Loop-D and β23/V5 regions of Env, conferred cross-resistance to several CD4bs bnAbs, while more potent CD4bs bnAbs like N6 and 1-18 largely retained their activity. Our findings demonstrate that prophylactic VRC01 can select for viral escape after infection, underscoring the need for next-generation bnAbs with improved breadth and potency to enhance durability and efficacy of antibody-based HIV prevention.

---

Broadly neutralizing antibodies (bnAbs) play a key role in both vaccine and passive immunization prevention strategies.^1,2^ The Antibody Mediated Prevention (AMP) trials, conducted in Africa (HVTN 703/HPTN 081) and the Americas/Switzerland (HVTN 704/HPTN 085), were landmark studies showing that passively administered bnAbs can prevent diagnosable acquisition of HIV.^3^ The AMP study found that CD4 binding site (CD4bs) antibody VRC01 was 75% effective against highly sensitive viruses (IC_80_ less than 1 µg/ml),^3^ and that serum neutralization titers were a reliable biomarker of protection.^4^ However, VRC01 efficacy declined with increased viral resistance, as well as greater genetic divergence in the VRC01 epitope relative to reference VRC01 sensitive strains.^3,5^ These findings underscore the challenge posed by HIV diversity and highlight the importance of understanding virus-bNAb interactions to inform both bNAb and vaccine studies.

In AMP, VRC01 transiently controlled viral load in individuals who acquired susceptible viruses, indicating antibody activity post-acquisition.^3,6^ This bnAb activity raised a concern that bNAb-based prevention strategies may select for viral resistance. This could limit the durability of the intervention, lead to the accumulation of resistance-conferring mutations among people who acquire HIV-1, reducing the effectiveness of future antibody-mediated interventions and vaccines. Although resistance has been documented in other pre-exposure prophylaxis (PreP) interventions,^7^ to date, *de novo* escape from bnAbs used for prevention has not been documented. In this study, we evaluated whether VRC01 selected for viral escape following HIV acquisition, and if so, whether this would compromise the effectiveness of other clinically relevant CD4 binding site bnAbs.

The determinants of viral escape depend on both viral factors, such as HIV diversity and fitness cost of a mutations, as well as host factors such as specificity and strength of the immune pressure.^8–10^ When used for prevention, bnAbs are present at the time of acquisition when viral diversity is typically very low thereby limiting the evolutionary pathways available for escape.^11–13^ In individuals treated in acute infection and later switched to VRC01 immunotherapy, no viral evolution was observed in rebound virus, suggesting a high barrier to escape in this setting.^14^ In contrast, during chronic infection, higher viral diversity provides more opportunities for resistance to emerge. VRC01 treatment often resulted in rapid viral rebound, due to either pre-existing resistant mutants or because the high diversity provides shorter evolutionary escape routes.^15–18^

Mapping VRC01 resistance is challenging due to the quaternary nature of the epitope and because escape mutations can occur in distal sites outside structurally defined contact sites.^19^ Resistance mutations have been identified in multiple regions of gp120, including Loop D, the CD4 binding loop, β20/β21, β23, and the V5 loop.^14,19–22^ Although bnAb resistance mutations provide a selective advantage when under antibody pressure, they are often found at a very low frequency among circulating viruses and have been linked to reduced viral entry efficiency and replicative capacity.^23–25^ Mutations that have a fitness cost may revert in the absence of selective pressure, although compensatory mutations can arise over time to restore fitness.^23,24,26^

In our studies of participants who acquired VRC01 susceptible viruses in the AMP trials, HIV deep sequencing revealed that escape mutations emerged post-acquisition in approximately a third of individuals (8/26) compared to none in the placebo group (0/21). These mutations conferred cross-resistance to some CD4bs bnAbs. However, the more potent generally remained effective. Resistant mutations were found at a very low frequency in global sequences. As rare amino acids can impose a fitness cost on the virus,^23–25,27^ it will be important to monitor their persistence after acquisition and their impact on fitness will be important going forward as an increase in resistant viruses would impact bnAbs-based interventions used in HIV prevention, treatment, and remission strategies.

## Results

### Identification of participants with single-lineage infections sensitive to VRC01 (IC_80_ less than 3 µg/ml)

In the AMP study, participants received infusions of saline (placebo) or 10 mg/kg (low-dose) or 30 mg/kg (high-dose) VRC01 every eight weeks, with HIV diagnostic testing every four weeks.^3^ Participants were grouped into (i) those diagnosed with HIV by the week 80 visit (i.e., within ~ 8 weeks of the last infusion) (trial endpoint); and (ii) those diagnosed with HIV after the week 80 visit (i.e., more than 8 weeks after the last infusion) (**Table 1)**. Infections were classified into single or multiple transmitted founder (TF) lineages based on deep sequencing of the env gene at the earliest RNA-positive plasma samples.^13,28^ Pseudoviruses representing the TF lineage and neutralization data linked to these lineages were available.^3^

**Table 1:** Number of participants who acquired single-lineage infections sensitive to VRC01, defined as IC_80_ < 3 µg/ml.

|  | HVTN 703/HPTN 081 (Africa Trial) |  |  |  | HVTN 704/HPTN 085 (Americas Trial) |  |  |  |
| --- | --- | --- | --- | --- | --- | --- | --- | --- |
|  | VRC01<br>10 mg/kg | VRC01<br>30 mg/kg | Placebo | Total | VRC01<br>10 mg/kg | VRC01<br>30 mg/kg | Placebo | Total |
| No. HIV-1 diagnosis* | 34 | 24 | 29 | 87 | 37 | 34 | 41 | 112 |
| No. single lineage<br>infections (Env)** | 22 | 11 | 17 | 50 | 19 | 26 | 29 | 74 |
| No. single VRC01<br>sensitive lineage | 9 | 4 | 10 | 23 | 6 | 7 | 11 | 24 |
| No. individuals with<br>variant VRC01 sites | 6 | 3 | 2 | 11 | 1 | 3 | 3 | 7 |
| No. env sequences at<br>time point 1 median<br>(range) | 33<br>(3-100) | 193<br>(71-245) | 123<br>(23-369) | 71<br>(3-369) | 155<br>(16-324) | 104<br>(3-379) | 181<br>(27-333) | 138<br>(3-379) |
| No. env sequences at<br>time point 2 median<br>(range) | 61<br>(5-184) | 81<br>(63-216) | 80<br>(4-208) | 74<br>(4-216) | 165<br>(30-192) | 112<br>(28-357) | 115<br>(42-467) | 116<br>(28-467) |

\*All participants who acquired HIV-1 were included except for 25 participants with sequence data from only a single time point.
\*\* Six participants were excluded because their first sequencing time point was $\geq 45$ days from the estimated date of acquisition (range 45-73 days).<sup>31</sup> Of the 47 single-lineage infections with sensitive viruses, 39 occurred by the week 80 visit (23 from HVTN 703; 24 from HVTN 704) and 8 occurred after the week 80 visit (2 from HVTN 703; 6 from HVTN 704).

This study examined participants who acquired a single transmitted founder lineage sensitive to VRC01 (IC_80_ <3 µg/ml), enabling us to explore how VRC01 pressure shaped viral evolution. Restricting the analysis to participants sampled within 45 days of estimated acquisition minimized the confounding effects of autologous CD4bs antibody responses, which typically emerge four to 16 weeks after acquisition.^29–31^

Forty-seven participants met criteria for inclusion (**Table 1)**: 23 in the Africa trial and 24 in the Americas trial. Of these, 39 were diagnosed by the week 80 visit, and 8 after the week 80 visit. A total of 12,008 env sequences were generated, with a median number per participant per time point of 108 (range 3 to 379). The median estimated date of acquisition of the first sequencing time point was 18 days (range: 7 – 45 days) and the intra-participant env diversity at the first sequencing time point was very low, consistent with recent acquisition (median DNA distance: 0.05%, range: 0.02 - 0.12%). The interval between sequencing time points 1 and 2 for the placebo group (trials pooled) was 14 days (range: 5 to 100 days) and for the VRC01 treatment group (dose group and trials pooled) was 12 days (range: 6 to 63 days).

### Evolution in the VRC01 epitope post-acquisition

To identify putative VRC01 escape mutations, we tracked amino acid changes over time at 34 amino acid sites within the VRC01 epitope **(Supplementary Table S1)**. Two complementary approaches were used. First, amino acid changes were visualized between time points using escape logograms, which compared time point 2 (TP2) sequences to the consensus sequence at time point 1 (TP1) **(Figure 1A, 1B; Supplementary Figure S1)**. Second, changes were mapped onto phylogenetic trees using highlighter plots to display amino acid amino acid changes from the transmitted founder lineage alongside the tree **(Figure 1C, 1D; Supplementary Figure S2)**. This allowed visualization of sequence evolution at different timepoints, as well as co-variation where changes were located on the same sequence.

**Figure 1.**
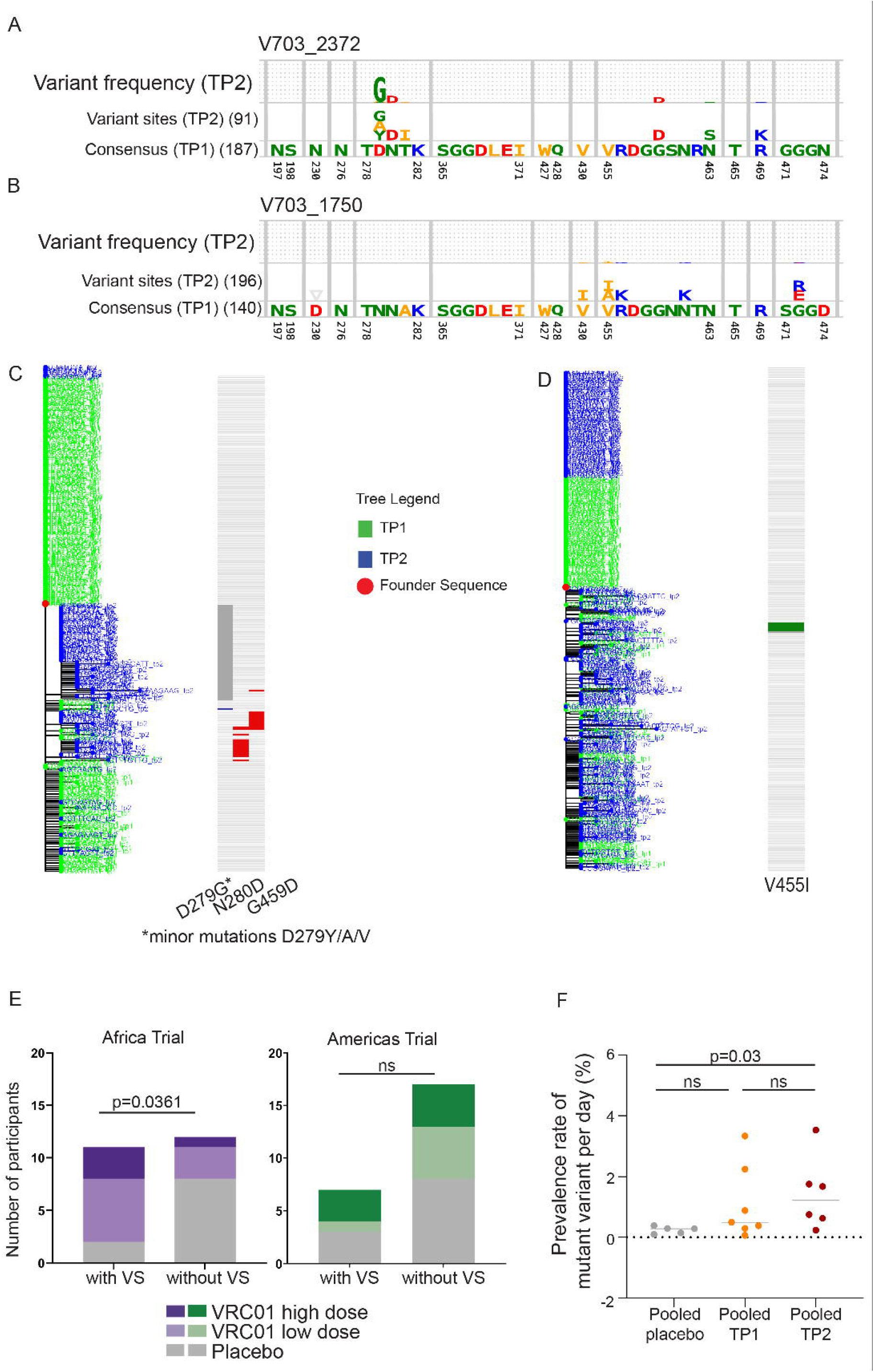
Identification of putative post-acquisition VRC01 escape mutations in individuals with single-lineage infections by VRC01-sensitive viruses (IC_80_ <3 µg/ml). A-D. Approach used to identify escape mutations within the 34 amino acid VRC01 epitope, comparing a VRC01 high-dose recipient (V703_2372, A and C) and a placebo recipient (V703_1750, B and D). **A and B**. Logoplots showing amino acid (AA) changes between time point 1 (TP1) and time point 2 (TP2). The bottom panel shows the TP1 founder lineage consensus, the top panel shows TP2 variant frequencies, and the middle panel is scaled to show variant AAs. Sequence counts are shown in parentheses. **C and D**. Phylogenetic trees of TP1 (green) and TP2 (blue) sequences. Adjacent highlighter plots display AA changes within the VRC01 epitope relative to the sequence closest to the TP1 consensus. **E**. Number of participants with VRC01 variant sites (VS) in the Africa Trial (purple) and Americas Trial (green). Compared to the placebo group, the treatment group (pooled) had more VS in the Africa trial (*p = 0.036 two-sided Fischer’s exact test), but not the America trial (p=ns). **F**. Change in VS per day pooled across trials and grouped by treatment: placebo (grey), VRC01 low dose (light brown) and VRC01 high-dose (burnt orange). A significant difference was observed between VRC01 high-dose group the placebo group (two-sided Mann-Whitney *p=0.030).

Variant sites present at >1% frequency of total sequences (time points pooled) were investigated further. Variant sites were identified in 18 of the 47 individuals, with a significantly higher frequency of individuals with variant sites in the VRC01 group in the Africa trial (groups pooled) compared to the placebo group 69% (95% CI: 42–87%); vs 20% (95% CI: 4–51%)(p=0.04, two-sided Fisher’s exact test), but not the Americas trial **(Table 1; Figure 1E)**. Furthermore, the change in frequency of variant sites between time points was significantly greater in the high-dose group compared to placebo based on change in frequency per day (trials and groups pooled) (p=0.03 by two-sided Mann-Whitney test) **(Figure 1F; Supplementary Figure S3)**.

Variant sites were most frequently detected in Env loop D at sites 276, 279, or 280, found in eight individuals and only in the VRC01 groups; 11 variant sites were observed only in the VRC01 group (N197T, D276T/I, D279G/A/Y/K, N280D, S365P, P369S/T, R456K, G458E, K460N, D462Y/G, D474N), whereas one variant site (E370K) was observed only in the placebo group, and three variant sites (T/V455I/A, G459E/K, and D461N/E) were observed in both the VRC01 and placebo groups.

### Single and double mutations that confer resistance to VRC01

Further investigation of participants with variant sites was conducted to determine whether these amino acid changes conferred VRC01 resistance. Variant sites were introduced into the corresponding VRC01 sensitive pseudovirus (PSV) clone representing the TF lineage identified in time point 1. The impact of the mutations on VRC01 sensitivity were assayed in the TZM-bl neutralization assay and defined as escape mutations if there was a more than three-fold change in IC_80_ **(Table S2; Supplementary Figure S4)**.

Escape mutations evolved post-acquisition in viruses of 31% (8/26; 95% CI: 17–50%) of individuals who received VRC01, but none in the placebo group (0/21), corresponding to a 95% CI odds ratio of >2.02 **(Figure 2A; Supplementary Table S2; Figure S5)**. All participants with VRC01 escape were diagnosed with HIV by the week 80 visit (within 8 weeks of the last VRC01 infusion), with no escape identified in participants who were diagnosed after the week 80 visit. Escape was more common in the Africa trial 46% (6/13, all subtype C), compared to the Americas trial 15% (2/13, all subtype B).

**Figure 2.**
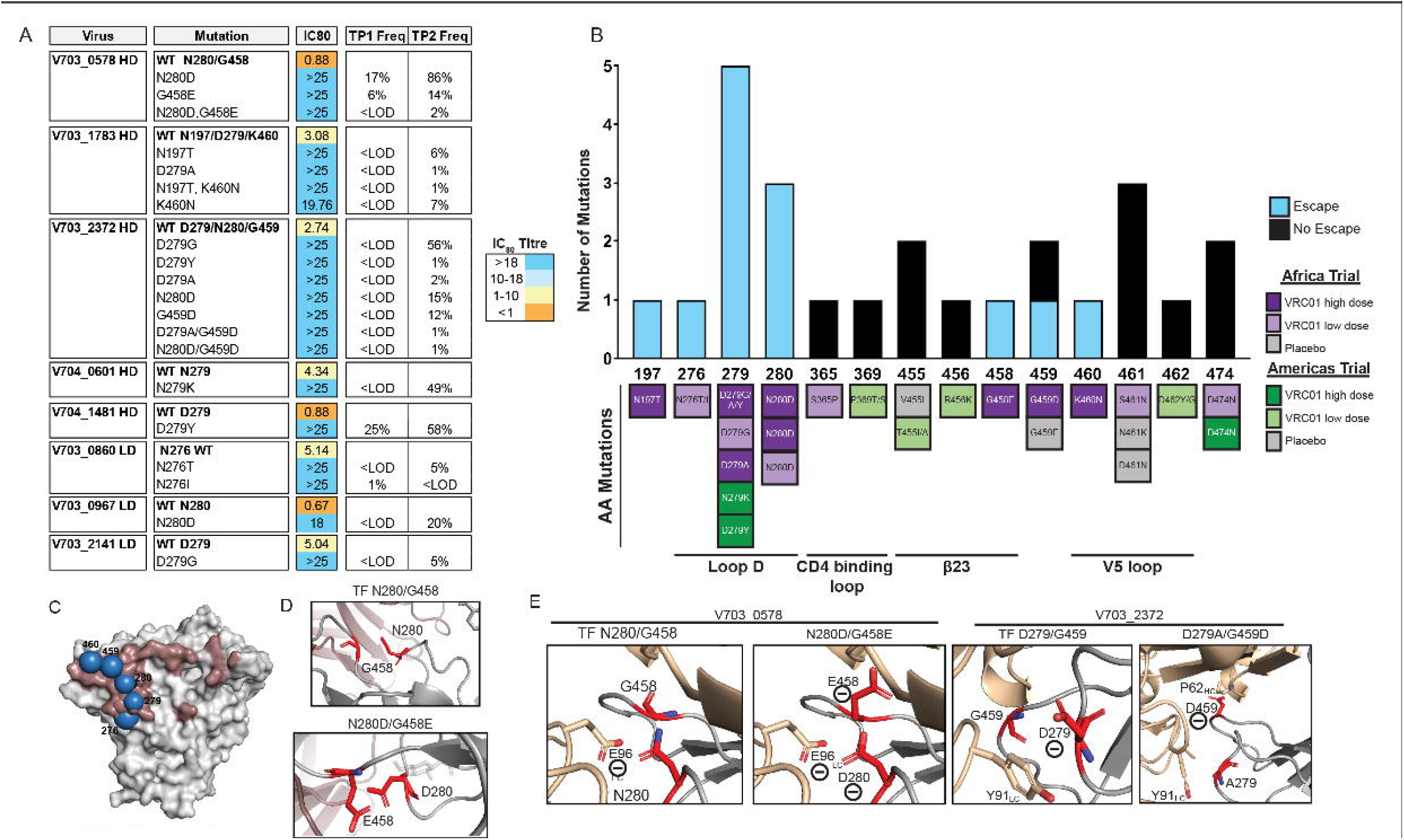
Post-acquisition VRC01 escape mutations. **A**. IC_80_ titers of the parental pseudovirus and the corresponding escape mutants in the VRC01 high-dose and low-dose groups. No escape mutations were detected in the placebo groups. The frequency of mutations at time point 1 (TP1) and TP2 is indicated. <LOD: below the limit of detection. **B**. Location of escape mutations on gp120. Some mutations conferred escape (more than 3-fold difference from TF in IC_80_, blue bars), and mutations that others had no effect on VRC01 sensitivity (less than 3-fold difference from TF in IC_80_, black bars). Mutations are colored by trial group. **C**. A crystal structure of the gp120 (PDB ID: 4LST) (grey), the VRC01 epitope footprint (red) and the escape mutations (blue). **D**. Close-up images of the interaction between VRC01 (wheat color) and the HIV gp120 protein (grey), with residues of interest shown in red. **E**. Close-up images showing interactions between VRC01 (wheat color) and gp120 (grey) with residues of interest are shown in red. Escape mutations were modelled using the mutagenesis tool in PyMOL Molecular Graphics System, version 2.4.2, Schrödinger, LLC.

The Loop D region was a hot spot for escape mutations, which were present in all eight participants having escape mutations at residues 276, 279, and/or 280. One participant had mutations at residue 276, five had mutations at residue 279, and three had mutations at residue 280 in three participants **(Figure 2A)**. One participant (V703_2372) had mutations at both residues 279 and 280. Toggling of resistant mutations was observed in two Loop D residues: N276T/I and D279G/Y/A in participants V703_0860 and V703_2372, respectively. Additional escape mutations were identified in the N197T glycan and the β23-V5 region at residues 458 (G458E), 459 (G459D), and 460 (K460N); with these mutations occurring in different participants **(Figure 2B)**. In a previous analysis of sequence features that impacted the efficacy of prevention in AMP, residue 460 was overall the most influential in the epitope distance analysis, with residue 459 being influential in the Americas trial.^5^

For six of the eight participants identified with escape mutations, the escape mutations were either below the level of detection or at a low frequency (<2%) at the first sequencing time point **(Figure 2A)**. Seven of the eight participants had sequence depths ranging from 107 to 379 **(Supplementary Table S2)**, providing 95% confidence for detecting variants present at frequencies greater than 0.8– 2.8%. In contrast, one participant (V703_0967) had only 42 sequences, corresponding to a 95% confidence threshold for detecting variants present at frequencies above 7%. In six of the eight participants, the final VRC01 infusion coincided with the day of HIV-1 diagnosis (day 0), and these high antibody concentrations during acute infection could have driven viral escape. The remaining two participants—one in the low-dose group (V703_2141) and one in the high-dose group (V704_1481)—received their last infusion 23 and 27 days before diagnosis, respectively (**Figure 3B-I)**.

**Figure 3:**
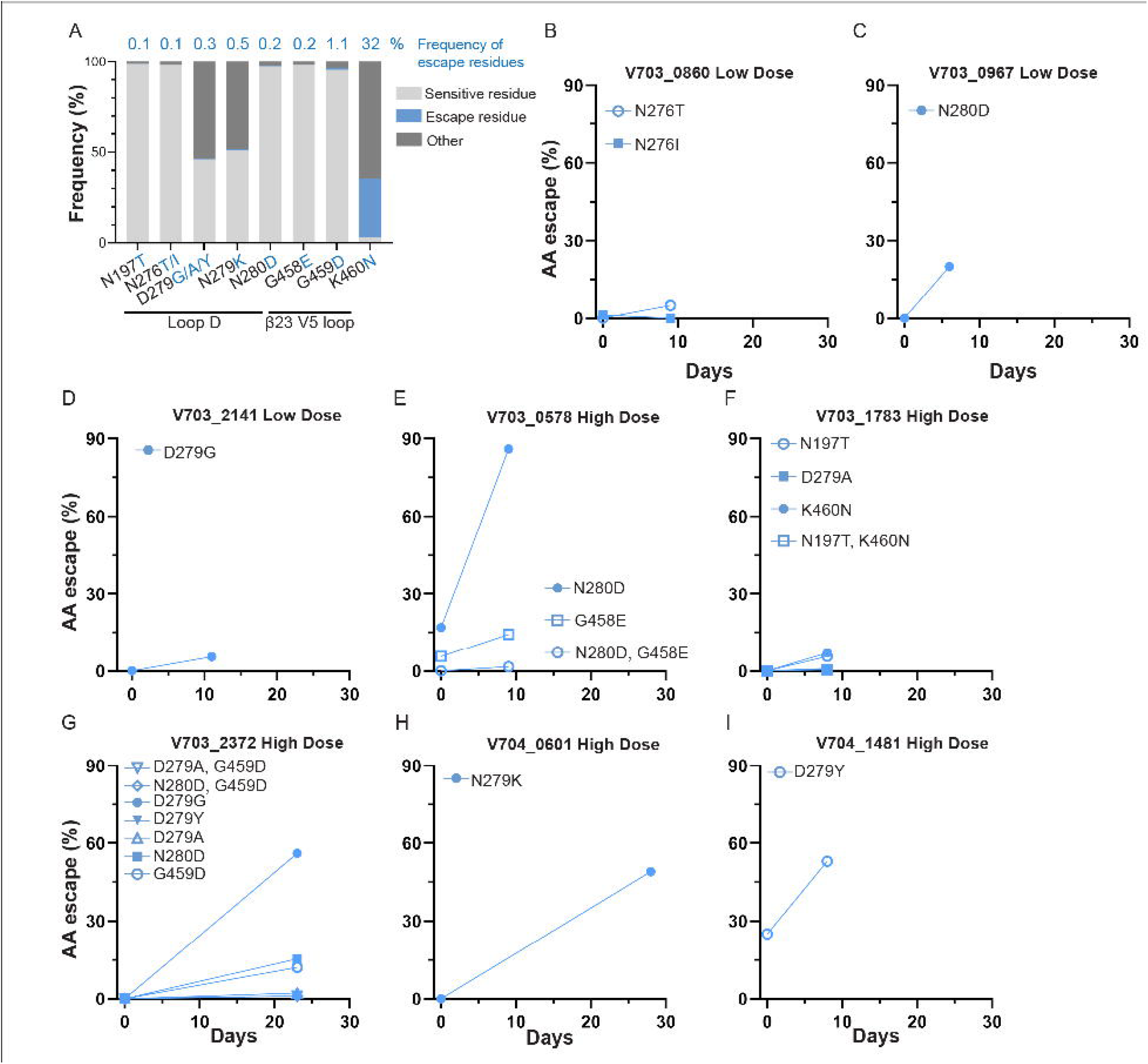
Frequency and kinetics of VRC01 escape following HIV diagnosis (Day 0). **A**. Database (8,452 filtered sequence alignment, https://www.hiv.lanl.gov/) frequencies of: wildtype residues (AA in the transmitted founder lineage, light grey), escape residues (blue), other (neither wildtype nor escape, dark grey). **B to I**. Frequency (%) at TP1 and TP2 of sequences harboring escape mutations. Two participants (V703_2141 and V704_1481) received their infusion 23 or 27 days prior to sequencing, while the remaining six received their last VRC01 infusion on the same day as sequencing (day 0). Number of sequences per time point is shown in Supplementary Table S2.

### Escape mutations are rare in global reference panels

Escape was identified at seven different VRC01 epitope positions across eight individuals. Most of these mutations were found at a frequency of less than 1% of the HIV sequences in the database (8,452 sequences, www.LANL.gov), except for K460N **(Figure 3A)**. The K460N change was a reversion from one relatively uncommon amino acid (11%) to a more common amino acid (32%) and was identified in one individual (V703_1783) whose viruses harboured multiple escape mutations. The amino acid frequencies of escape mutations in subtype-matched reference panels were similar to the global panel with all mutations found below 1%, except for K460N which was found in 23% of subtype C viruses and 29% of subtype B viruses **(Supplementary Figure S4)**.

### Mechanisms of escape of single and double mutations

The mechanisms of immune escape were further explored by modelling the key mutations on a crystal structure of VRC01 bound to a subtype C gp120 protein, selected to match the subtype of the sequences from the participant **(Figure 2C)**. In this study, we identified three double mutations spanning loop D and the V5 loop—N280D/G458E, D279A/G459D, and N280D/G459D. These residues are close to each other on the envelope trimer **(Figure 2C, 2D)**. The N280 sensitive residue is polar but uncharged residue, and the D280 resistant mutation introduces a negative charge, which would result in repulsion from the nearby negatively charged E96 residue on the VRC01 light chain. The sensitive residues G458 and G459 are compact, nonpolar amino acids and the change to resistant mutations E458 or D458 introduces negatively charged amino acids with longer side chains. These combined mutations likely facilitate antibody escape by changing the nature of the epitope-paratope interaction. An alternative non-charge-dependent mechanism involved changes in side-chain size. For example, mutation of D279 to either A279 or G279 results in shorter side chains in addition to removal of the negative charge **(Figure 2D)**. Thus, modeling suggests that coordinated mutations in critical loops reshape the viral surface, enabling HIV to evade neutralization through both charge-based and other mechanisms affecting conformation.

### Some clinically relevant CD4bs bnAbs are unaffected by VRC01 escape

We screened seventeen of the twenty PSV mutants that conferred VRC01 resistance against other VRC01-class CD4bs bnAbs (VRC01.23LS, VRC07-523LS, 3BNC117, N6, VRC_CH31),^32–34^ as well as non-VRC01 class CD4bs bnAbs (1-18, CH235.12).^21,35^ As controls, we included PG9 and 10E8v4, which target the V2 apex and MPER, and not expected to be impacted by VRC01 escape.

In a pairwise correlation heatmap, the antibodies segregated into two main clusters (**Figure 4A)**. The first cluster consisted of VRC01-class bnAbs, whose titers strongly correlated with VRC01 titers (range 0.49-0.96, p=1.2×10^−15^ – 3×10^−7^ by ANOVA test; **Figure 4B, Supplementary Table S3**). In both the hierarchical clustering and correlation heatmaps **(Figure 4A, 4B)**, 3BNC117 and VRC_CH31 clustered with VRC01, with VRC01.23LS and VRC07-523LS forming a separate subcluster. N6 and 1-18 clustered separately. N6 titers only moderately correlated with VRC01 (coefficient of 0.32, p=0.02, **Fig. 4B, Supplementary Table S3**), whereas 1-18 and CH235.12 showed no significant correlation, nor did the control bnAbs PG9 and 10E8v4. Overall, most of the VRC01-class bnAbs overlapped significantly in mutations that affect escape, consistent with overlap in their epitopes. In contrast, N6 and 1-18 were largely unaffected by VRC01 escape, despite also targeting the CD4bs.^21,34^

**Figure 4.**
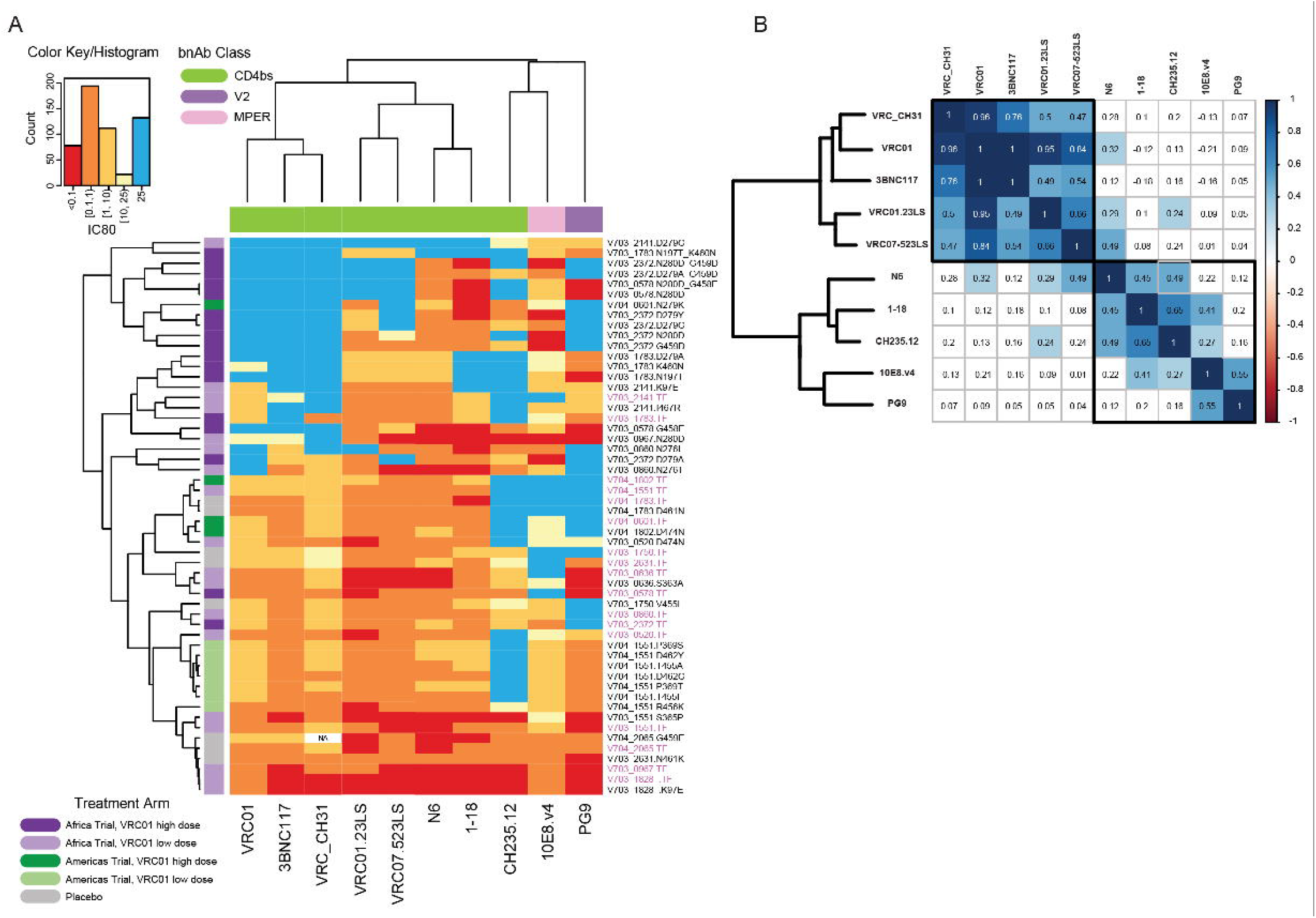
Relationship between escape mutations and bnAb sensitivity. **A**. Heatmap of IC_80_ neutralization titers of 54 pseudoviruses (rows) against 10 bnAbs (columns), 8 targeting the CD4bs epitope (labelled in green at the top), two controls [bnAb targeting the V2 apex epitope (lavender) and the MPER (pink)]. Rows and columns are ordered using hierarchical clustering: the top dendrogram illustrates the clustering of the bnAbs, whereas individual viruses are listed on the right hand side, with TFL plasmids in pink and mutants in black. The left side dendrogram illustrates the grouping of viruses based on their neutralization resistance levels, with the most sensitive viruses clustering at the bottom of the heatmap (dark red and orange hues) and the more resistant ones at the top (light yellow and blue). Colored bands on the left indicate trial study group (Treatment arm). **B**. Heatmap showing all pairwise correlation coefficients across all pairs of bnAb neutralization titers (columns in the heatmap shown in 4A). Correlation coefficients were calculated using random effects generalized linear models (GLMs) accounting for the Env backbone as a random effect. White cells indicate no statistical significance, whereas statistically significant correlation coefficients are by colored cells, color coded from red (perfect negative correlation) to dark blue (perfect positive correlation).

We next assessed whether the hierarchical clustering of viruses with VRC01 escape mutations corresponded to the observed reduction in potency of CD4bs-directed antibodies. **Figure 5** shows IC_80_ titers of the bnAbs against the parental pseudoviruses **(5A)** and mutated pseudoviruses **(5B)**, with the pie charts summarising their IC_80_ profiles. Viruses sensitive to VRC01 were generally sensitive to other VRC01-class antibodies, with VRC07-523LS, VRC01.23LS, N6 having significant increases in potency compared to VRC01 (FDR q<0.01 by paired Wilcoxon test after multiple testing correction). **Figure 5B** shows the IC_80_ titers for the same antibodies against the viruses carrying VRC01 escape mutations. Here, we excluded PSVs where the parental PSV was resistant to specific antibodies including: VRC-CH31 (1 virus), 3BNC117 (2 viruses), or CH235.12 (3 viruses). VRC01 escape mutations had similar effect on potency of VRC01, VRC-CH31 and 3BNC117, however they did not significantly affect the potency of VRC07-523LS, VRC01.23LS and N6). bNAb 1-18 showed the greatest resilience, retaining activity with IC_80_ values below 1 µg/ml against 75% of VRC01 escape variants.

**Figure 5.**
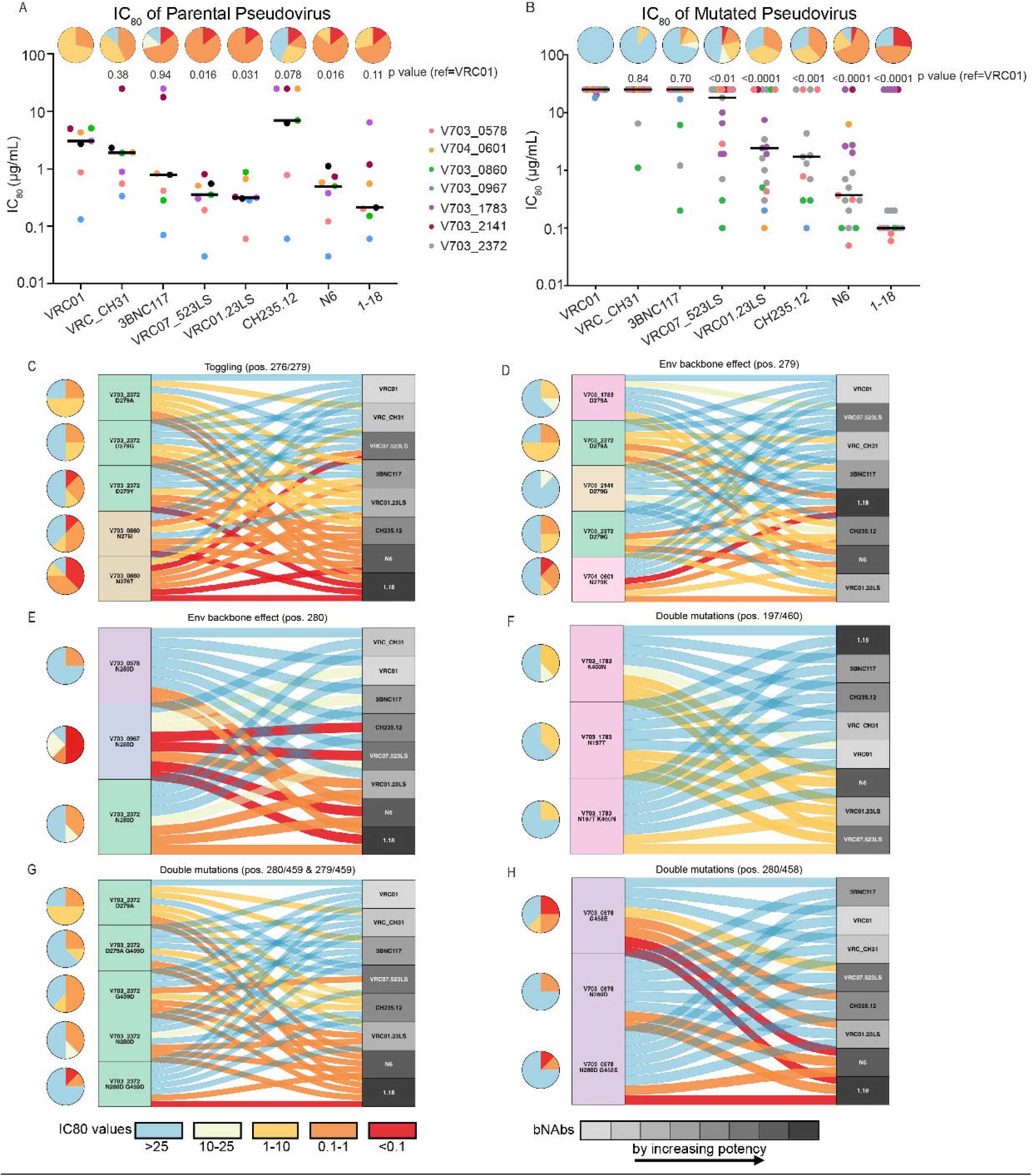
IC_80_ titers of parental sensitive pseudoviruses (PSV) and corresponding resistant clones to bnAb panel. Pie charts (A-H) showing resistance profiles IC_80_ (µg/ml), with color key at the bottom of panel G. **A-B**. Scatter plot showing parental PSV IC80 values against different bnAbs **(A)** and the effect of VRC01 escape mutations on each bnAb sensitivity (IC_80_) **(B)**. Bars indicate the median IC_80_ (µg/ml), and 2-way paired Wilcoxon p-values denote whether they were statistically different compared to VRC01. **C - H**. Effect of single mutations **(C**,**D**,**E)**, in different parental PSV backbones **(D**,**E)**, and double mutations combinations **(F, G, H)** on bnAb cross-recognition. The alluvial flow links mutant PSV sensitivity (colored by IC_80_ value) (left) to bnAb (right). The bnAbs are colored by potency (right), with the darker shades of grey showing the increase of bnAb IC_80_/ potency as determine in A. The potency is ordered differently on each plot to reflect differential impact of the different mutations. All amino acid positions correspond to HXB2 reference HIV virus.

The alluvial plots **(Figure 5 C to H)** illustrate the relationship between the escape mutations (left side) and the bnAb (right side). Each plot shows pie charts summarizing the IC_80_ profile of a given mutant virus and colored flows (alluvia) connecting each mutant to the bnAb, colored by IC_80_ titer. The bnAbs are shaded from light to dark grey based on their overall potency against the escape variants identified in **Figure 5B**.

The Loop D mutations were most frequently observed **(Figure 2B)**, but their impact on resistance varied depending on both the specific mutation and the Env backbone in which they occurred **(Figure 5C, D, E)**. Furthermore, at positions 276 and 279, alternative amino acids (i.e., toggling escape mutations) within the same backbone had different effects on resistance. For example, in the V703_0860 backbone, N276I conferred greater resistance than did N276T. Similarly, in the V703_2372 backbone, D279G or Y conferred greater resistance than did D279A (**Figure 5C)**. The impact of D279A, D279G and N280D were also context dependent **(Figure 5D and E)** – the N280D mutation conferred resistance to 6/8 bnAbs in the V703_0578 backbone, compared to 4/8 bnAbs in the V703_2372 backbone and only 1/8 bnAbs in the V703_0967 backbone **(Figure 5E)**. Similar backbone effects were observed for the D279A and D279G mutations.

When double mutations were observed, the relative contribution of each mutation varied. In V703_1783, the N197T/N460K double mutant conferred greater resistance to CD4bs antibodies than either mutation tested alone **(Figure 5F)**. Similarly, in V703_2372, both double mutants (N280D/G259D) and (D279A/G249D) showed increased resistance compared to the corresponding single mutants identified in that participant **(Figure 5H)**. In contrast, the V703_0579 N280D/G458E double mutant showed no greater resistance than the N280D single mutant (both were resistant to 6/8 bnAbs) **(Figure 5G)**.

Regardless of the mutation or backbone, the next generation bnAbs 1-18 and N6 were generally resilient to escape, as indicated by their position towards the bottom of the alluvial plots in five out of six cases.

## Discussion

The Antibody Mediated Prevention (AMP) trials were the first demonstration that a single broadly neutralizing antibody (bnAb), VRC01, could prevent HIV acquisition by sensitive strains._3_ These trials provided an opportunity to explore viral evolution in the setting of antibody pressure immediately following acquisition. Our findings showed the emergence of VRC01 escape mutations post-acquisition in approximately one-third of individuals who acquired sensitive viruses. These mutations reduced the effectiveness of some CD4bs bnAbs, but had a limited effect on more potent antibodies such as N6 and 1-18. The rarity of these escape mutations in circulating viruses suggests they impose a fitness cost, and as VRC01 pressure is transient, there may be limited opportunity for compensatory mutations to arise and restore fitness.^23–25^ This raises uncertainty about whether resistant variants will persist in these individuals where onward transmission could undermine future antibody-based prevention and therapeutic interventions.

In two cases, escape occurred prior to diagnosis, likely due to low in vivo VRCO1 titers in the last four weeks post infusion which allowed ongoing replication.^4,36^ However, VRC01 infusions were administered at the visit at which HIV diagnosis was subsequently made in six cases, and of these, five did not have detectable escape at the time of diagnosis. In these cases, the high VRC01 levels during the exponential viral expansion during acute infection may have played a role in selecting for escape. However, it is also possible that escape variants were present at a very low frequency at diagnosis and only became detectable later. These observations highlight the need to maintain high therapeutic antibody levels, ideally through combination strategies that target multiple epitopes simultaneously.^18,37,38^

We found that a single escape mutation was sufficient to confer VRC01 resistance with most escape mutations clustered in the Loop D region of the Env protein, an important contact site for VRC01 and other VRC01-class antibodies.^16,19,21,22,34,39^ However, some viruses acquired two or more escape mutations at different sites spanning loop D and V5. Structural modeling suggests that double mutations may altering the amino acid side chain and increasing the negative charge of the epitope, each of which could repel VRC01 binding. We were unable to confirm this hypothesis in neutralization assays since the level would have been above the maximum VRC01 concentration tested (25 µg/ml). We did observe, however, cases of viruses with double mutations conferring greater cross-resistance to other bnAbs than conferred by single mutations in the same backbone (**Figure 5, F to H)**. For example, in participant V703_2372, PSV with the single G459D, N280D and D279N mutations were neutralized by VRC01.23LS, however, the combination of G459D mutation with either N280D or D279N conferred cross-resistance (**Supplementary Figure S5)**.

All CD4bs antibodies evaluated in this study belonged to the VRC01-class, except for 1-18 and CH235. Antibodies in this class share a common heavy chain and mode of binding, and while there is epitope overlap within a class, they all have unique features accounting for their differences in potency and breadth.^19,21,22,34,35,40^ Our results showed that while VRC01 escape mutations conferred cross-resistance to other VRC01 class antibodies, N6 was less affected. This reduced impact on N6 bnAbs can be attributed to a different binding angle (5-8 degrees different), which results in a 20% larger loop D binding surface and less steric hindrance compared to other VRC01-class antibodies, thus allowing it to accommodate more variation in the V5 loop.^34^

The bnAb 1-18 is a non-VRC01-class CD4bs antibody with exceptional potency and breadth, neutralizing 97% of viruses in a multiclade panel and 90% in a clade C panel, with a geometric mean IC_80_ of 0.183 µg/ml and 0.279 µg/ml, respectively.^41^ Its enhanced potency has been attributed to improved ability to tolerate glycans in the Loop D region. While this bnAb was largely resilient to VRC01 escape mutations, mutations at position 279 affected sensitivity to 1-18 in some backbones: the D279G in the V703_2141 backbone mutation increased the resistance by 20-fold but had no effect in the V703_2372 backbone. The V703_1783 backbone was sensitive to bnAb 1-18 (IC80 0.37 ug/ml), and while the introduction of N197T and K460N alone increased resistance by ~7 fold (IC80 2.0 to 2.57 ug/ml), the combination of these mutations in this backbone increased resistance by 68-fold (IC80 >25 ug/ml) (**Supplementary S4)**. While these specific changes have not been reported to affect bnAb 1-18 neutralization, they are in the antibody contact sites.^21^ Overall, bnAb 1-18 retained its potency with a median IC_80_ < 0.1 µg/ml against VRC01 escaped viruses.

This study found that HIV escape mutations against VRC01 that arise soon after acquisition do not differ from those identified in people living with HIV (PLWH) treated with VRC01 during chronic infection. However, we did not assess distal sites outside the VRC01 epitope that may affect sensitivity.^19,39^ We also did not investigate multiple lineage infections where escape may be facilitated through recombination, particularly in individuals who acquire viruses with different VRC01 phenotypes.^42,43^

Escape from VRC01 was predominantly located in the D loop of the HIV envelope protein, a critical region recognized by CD4bs bnAbs and a focus of prevention, treatment and remission (functional cure) trials.^1,15,33,44^ The CD4 binding site is also important target in vaccine development where most germline-targeting strategies aim to elicit VRC01-like bNAb responses, with early vaccine trials having yielded encouraging results.^45–47^ Understanding the HIV mutational landscape within this region provides insights into the boundaries of antibody recognition and highlight the specific vulnerabilities that may undermine protection. These data underscore the need to account for natural escape pathways within immunogen design, as well as bNAb combinations, aimed at durable escape resistance.

This study provides new insights into how bnAbs used in prevention shape viral evolution during the earliest stages of infection. By using deep sequencing data, we captured key early evolutionary events and identified resistance mutations that conventional sequencing methods would have missed. Our findings also highlight the risk of resistance arising due to incomplete neutralization, a challenge currently being addressed by combining bnAbs targeting multiple epitopes.^1,48–52^ We found that VRC01 escape mutations affected CD4bs bnAbs differently, suggesting that combining antibodies within the same class could also be beneficial, particularly if escape forces the virus down a vulnerable evolutionary path. As newer bnAbs with greater potency, broader coverage, and longer half-lives enter clinical evaluation, our data provide a framework for predicting and mitigating viral resistance. Together, these advances provide insight that informs the development of antibody-based prevention strategies aimed at resilience against HIV evolution.

## Methods

### Ethics Statement

All participants in the AMP trials provided written informed consent. The work at the University of Cape Town was approved by the UCT Human Research Ethics Committee (HREF ref. no. 176/2017), and the work at National Institute of Communicable Diseases was approved by the University of the Witwatersrand Human Research Ethics Committee (protocol no. M201105). The TZM-bl neutralization assay studies at Duke were approved by the Duke University Health System Institutional Review Board (Duke University, protocol no.Pro00093087). Viral sequencing at the University of Washington was deemed not to require Human Subjects approval by the Institutional Review Board.

### Study populations

See AMP study.^3^ In brief, HVTN 703/HPTN 081 and HVTN 704/HPTN 085 were two parallel phase 2b, multicenter, randomized, double-blind, placebo-controlled, proof-of-concept efficacy trials of the broadly neutralizing HIV monoclonal antibody (bnAb) VRC01 in women with a high likelihood of HIV acquisition (HVTN 703/HPTN 081, conducted in sub-Saharan Africa referred to as the Africa Trial) and cisgender men and transgender persons with a high likelihood of HIV acquisition (HVTN 704/HPTN 085, conducted in North America, South America, and Switzerland referred to as the Americas Trial).^3^ Participants were healthy adults without HIV who provided written informed consent after an evaluation of their understanding of the protocol. HVTN 704/HPTN 085 enrolled adults aged 18 to 50 years. HVTN 703/HPTN 081 enrolled adults aged 18 to 40 years.

Participants received either 10 mg per kilogram (low dose, LD), 30 mg per kilogram (high dose, HD) or saline placebo every 8 weeks and were tested for HIV infection monthly. The earliest RNA-positive plasma sample was sequenced and the dominant infecting viruses (>10% of the population) were evaluated for VRC01 neutralization sensitivity using the TZM-bl assay.^3^ Viral sequences were also obtained from a second sampling visit 1-3 weeks later. The calculation of estimated dates of diagnosable acquisition (EDDA) were provided by Rosenthal et al.^31^ In summary, the EDDA was estimated using a Bayesian posterior distribution, which combined independent inputs from diagnostic timing estimators with combined gag-Δpol and *rev-env-Δnef* based estimators utilizing Poisson Fitter 2.0.^31,53,54^ In this study, the date of acquisition was assumed to be 7 days prior to the EDDA to account for the infection eclipse phase.^55^

### Sequencing

HIV-1 was sequenced using the Pacific Biosciences single-molecule real-time SMRT platform from amplicons derived from single genome amplification (SGA)(3) tagged with unique index adaptors for sequencing or from a UMI-based approach.^56^ For the UMI–based approach, each viral genome was tagged with a unique molecular identifier (UMI) during cDNA synthesis, unused cDNA primers were removed using magnetic beads (Beckman Coulter TM Agencourt RNAClean XP), and the rev, env, nef (REN) genomic region (~3 kb; HXB2 position 5970 – 9012) was amplified. To minimize recombination during PCR, a limited number of templates were amplified per PCR reaction, estimated by endpoint dilution, such that no more than 25 amplifiable copies of cDNA were added to each reaction (8 repeats). The number of amplifiable templates was estimated using the Quality tool^57^ (ttps://quality.fredhutch.org/). The first round PCR products were size-purified using the Blue Pippin instrument (Sage Science), followed by a second round of PCR. For sequencing, SMRT bell adaptors were ligated to the PCR products, and the amplicons were sequenced on the PacBio platform using Sequel instruments. Circular consensus sequences (CCS) were processed into template specific consensus sequences using custom bioinformatics pipelines, for SGA (https://github.com/MullinsLab/sga_index_consensus) or UMI-containing samples (http://github.com/MurrellGroup/PORPIDpipeline). Briefly, CCS reads were filtered for length and quality, then demultiplexed to generate consensus sequences linked to a unique index or sample ID and UMI. Sequences were checked for contamination against a laboratory database before nucleotide and amino acid alignment using MUSCLE algorithm (https://www.ebi.ac.uk/jdispatcher/msa/muscle). Identical sequences were collapsed for the alignment review, and three rounds of independent review were performed.

### Viral characterization for participant selection

For the purposes of this study, we only included participants infected by a single lineage that was sensitive to VRC01 (IC_80_ ≤3.5 µg/ml).^3,5^ Lineage assignment is described in detail by Mullins et al.^28^ Briefly, two major approaches were taken: the poisson fitter method applied to the first sequencing time point only,^53^ and a phylogenetic/distance method which analyzed data from all timepoints. The distance/phylogenetic approach generated maximum-likelihood phylogenetic trees from all timepoints pooled. Nucleotide changes from the most common sequence at the first time point were visualized using highlighter and matcher plots in Phylobook (https://github.com/MullinsLab/phylobook).^58^ Trees with single clades with low pairwise distances were classified as single lineage infections, while multiple lineage infections had more than one clade on the phylogenetic trees with intra-clade distances, after exclusion of recombinant and hypermutated sequences, less than the inter-clade distances.

### Identification of amino acid variant sites in the VRC01 epitope

We defined the VRC01 epitope as amino sites that were VRC01 contact sites as well as having supporting *in vitro* evidence demonstrating VRC01 escape (**Supplementary Table S1)**. We also included site 230, which was associated with prevention efficacy in the Americas trial in virus sequence sieve analysis^5^. These sites were in the following domains: glycosylation (197,230); CD4 contact (198;459;472-474); loop D (276,278-281); CD4 Binding loop (365-371); β20,β21 / CD4 contacts 427-428,430; β23/CD4 contacts (456-458); V5 loop (460); loop β24 (465,469, 471).

All sites were numbered according to HXB2 coordinates. Intra-participant differences from the VRC01 epitope were visualized using a tool called Mupundu (https://github.com/HIVDiversity/Mupundu_public/tree/main?tab=readme-ov-file). Mupundu is written in R and used the Shiny package and the ggtree package. It takes three input files: a fasta file flagged by time point, a tree file in newick format and a file containing the information about the site of interest (site position, amino acids associated with resistance/sensitivity). From these inputs, Mupundu displays a tree alongside a highlighter plot, showing differences in the VRC01 epitope compared to the sensitive pseudovirus sequence in time point 1 (putative transmitted founder sequence). Amino acid residue positions were classified as variant sites (VS) if there were 5 or more viral sequences with an amino acid change at a particular site when compared to the transmitted founder lineage consensus sequence in both time points.

### Mutagenesis

Variant sites were introduced into the matched participant founder virus envelope using site-directed mutagenesis (Agilent QuikChange Lightning or QuikChange Lightning Multi Site-Directed Mutagenesis Kits) as per the manufacturer’s instructions. Mutations were confirmed with Sanger sequencing.

### Pseudovirus production and Neutralization assays

Pseudoviruses were produced from the founder and mutant plasmids, and VRC01 sensitivity directly compared using the TZM-bl neutralization assay. For the generation of HIV-1 Env-pseudotyped virus, 293T/17 cells were co-transfected with *env* plasmids and an *env*-deleted backbone vector (pSG3delEnv). The pseudovirus representing the TF lineage and mutant pseudoviruses were then titrated to determine the optimal dilutions to use in the TZM-bl neutralization assay.^59^ Neutralization sensitivity of the viruses to a panel of bnAbs was tested in the TZM-bl neutralization assay. Briefly, neutralization was measured as a function of reduction in luciferase reporter gene expression (due to the presence of bnAbs) after a single round of HIV-1 Env-pseudotyped virus infection of TZM-bl target cells. Neutralization titers are expressed as the concentration of the VRC01 drug product at which RLU were reduced by either 50% (IC_50_) or 80% (IC_80_) relative to virus control wells after subtraction of background RLU in cell control wells.

### Frequency of escape sites

To determine frequency of amino acid sites conferring escape in global virus panels, Env amino acid alignments of 8,452 group M HIV-1 sequences were downloaded from the Los Alamos HIV sequence database (http://ww.hiv.lanl.gov). This alignment contained only one sequence per participant, with very similar sequences and problematic sequences deleted. For frequencies in HIV subtype B and subtype C a subset of 3,043 and 1779 sequences were extracted respectively.

### Structural modelling

To visualize the location of the escape mutations of interest we used a crystal structure of the HIV Env gp120 protein in complex with VRC01 (HIV-1 subtype C strain ZM176.66 PDB ID: 4LST). The VRC01 antibody footprint was mapped by determining all residues on the gp120 protein that were within 4Å or either the heavy or light chain of the antibody. The escape mutations were modelled and the interactions they make with the VRC01 mAb were compared to the original residues at the same location. All analyses were conducted in Pymol v2.5.2.

### Statistical Analysis

The comparisons of participant numbers with variant sites between trial and treatment arms were calculated with two-sided Fisher’s exact tests performed using Graphpad Prism version 10 (Graphpad Software, USA). The neutralization titer heatmap with hierarchical clustering and the pairwise correlation heatmap were generated using the R statistical platform, version 4.2.1. Correlation coefficients between bnAb pairs were calculated using random effects generalized linear models (GLMs) for each bnAb pair using IC_80_ titers as fixed effect and env backbone as random effect. Significance of each GLM coefficient was calculated using the ANOVA test. Statistical comparisons of the IC_80_ titers of the WT viruses between VRC01 and all other bnAbs were conducted using 2-way paired Wilcoxon tests, and FDR multiple testing correction was applied with a threshold of q<0.1.

## Data availability

The GenBank accession numbers for the HIV Env clones used in the TZM-bl target cell neutralization assay are: HVTN 703/HPTN 081 sequences, ON890939–ON891092; HVTN 704/HPTN 085 sequences, ON980814–ON980967. The GenBank accession number for the AMP sequences are ON890939–ON891092; ON980814–ON980967; PX890939-PX026359; ON890939-ON891092; PX184492-PX150096 ON980967-ON980814; OR508960-OR508936; OQ912897-OQ912888.1. The following computations tools were used: PORPIDpipeline: https://github.com/MurrellGroup/PORPIDpipeline.git;

Sga_index_consensus: https://github.com/MullinsLab/sga_index_consensus;

EscapeLogos: https://github.com/HughMurrell/EscapeLogos.git

## Author contributions

Conceptualized the study and interpreted findings: CW, PLM, JIM, NM, LC, EEG, RR, PTE, LM; collected and analyzed the data: CW, PM, JIM, NM, LC, EEG, TB, BL AAM, HM, CAM, RR, PTE, LM; wrote original draft and revised subsequent versions of the manuscript: CW, JIM, PLM, NM, LC, EEG, TB, NMGG, RR, PTE; reviewed and edited previous versions. CW, LC, NM, JIM, PLM, EEG, NMGG, CAM, STK, MJ, RR, PTE, LM; all authors read and approved the final manuscript. LC, PBG, CW provided funding.

## Acknowledgments

We gratefully acknowledge the HVTN 703/HPTN 081 and HVTN 704/HPTN 085 study participants, the protocol development and study implementation teams, and the NIH/NIAID Vaccine Research Center for the clinical development and manufacturing of the study product.

## Supplementary information

Supplementary Figures and Tables are provided in a separate document.

## Supplementary material

**Table S1**. VRC01 Epitope amino acid positions

**Table S2**. Metadata of participants with variant sites

**Table S3**. Correlation coefficients calculated from random effects generalized linear model of Envelope backbones

**Figure S1**. Escape logograms for participants with variant sites in the 34 VRC01 epitope sites of interest. HVTN 703/HPTN 081: A-K; HVTN704/HPTN 085 L-Q. Logograms showing amino acid (AA) changes between time point 1 (TP1) and time point 2 (TP2). The bottom panel presents the founder lineage consensus sequence at TP1, while the top panel shows the frequencies of variants at TP2. The middle panel highlights variant amino acids identified at a site compared to the consensus at TP1 and has been scaled for visibility. Sequence counts at each time point are shown in parentheses.

**Figure S2**. Phylogenetic trees (left) displaying sequences from time point 1 (green) and time point 2 (blue). Adjacent highlighter plots show amino acid changes in the VRC01 epitope sites relative to a viral sequence (red dot) closest to the consensus at time point 1.

**Figure S3**. A graph indicating the change of prevalence of viral sequences with variant sites between sample time points. The median number of days between sampling time points: Placebo = 8 days (range 5-28), VRC01 low dose = 11 days (range 6-63) and VRC01 high dose =16 days(range 8-28).

**Figure S4: Heat plot of IC**_**80**_ **neutralization titers of pseudoviruses and corresponding mutants**. Plot on the lefthand side illustrating the IC_80_ titers, and the plot on the right-hand side showing the fold difference between the sensitive PSV and the mutant. The eight CD4bs bnAb shown on the top, together with two control bnAbs targeting V2 apex and MPER (PG9 and 10E8v4)

**Figure S5**. Frequencies of wildtype residues (AA in the transmitted founder lineage, light grey), escape residues (blue), other (neither wildtype nor escape, dark grey). A. All sequences (n=8,452); B. Subtype C sequences (n=1779); C. Subtype B sequences (n=3043). Filtered sequence alignments downloaded from the Los Alamos database (https://www.hiv.lanl.gov/)

