## Supplementary Tables for "VRC01 Selects Rare HIV Escape Mutations After Acquisition in Antibody-Mediated Prevention Trials"

**Table S1.** List of 34 amino acid positions/sites within the VRC01 epitope that were evaluated for changes over time.

| HXB2 amino acid position/site | Feature | Envelope Domain | Experiment | References/Citations |
|---|---|---|---|---|
| 197 | Glycosylation site | Glycosylation site | Neutralization | 1 |
| 198 | Glycosylation site | CD4 contact | Neutralization | 1 |
| 230 | Glycosylation site | Glycosylation site, Sieve site | Sieve analysis | 2 |
| 276 | VRC01 contact site | Loop D | Neutralization | 3 |
| 278 | VRC01 contact site | Loop D | Neutralization | 4 |
| 279 | VRC01 contact site | Loop D (CD4 contact) | Neutralization | 3 |
| 280 | VRC01 contact site | Loop D (CD4 contact) | Neutralization | 4 |
| 281 | VRC01 contact site | Loop D (CD4 contact) | Neutralization | 4 |
| 282 | VRC01 contact site | Loop D (CD4 contact) | Neutralization | 6 |
| 365 | VRC01 contact site | CD4 Binding loop | Neutralization | 7 |
| 366 | VRC01 contact site | CD4 Binding loop | Neutralization | 6 |
| 367 | VRC01 contact site | CD4 Binding loop | Neutralization | 6 |
| 368 | VRC01 contact site | CD4 Binding loop | Neutralization | 6 |
| 369 | VRC01 contact site | CD4 Binding loop | Neutralization | 1 |
| 370 | VRC01 contact site | CD4 Binding loop | Neutralization | 6 |
| 371 | VRC01 contact site | CD4 Binding loop | Neutralization | 6 |
| 427 | VRC01 contact site | β20, β21 (CD4 contacts) | Binding | 1 |
| 428 | VRC01 contact site | β20, β21 (CD4 contacts) | Binding | 1 |
| 430 | VRC01 contact site | β20, β21 (CD4 contacts) | Binding | 1 |
| 455 | VRC01 contact site | β23 (CD4 contacts) | Neutralization | 7 |
| 456 | VRC01 contact site | β23 (CD4 contacts) | Neutralization | 8 |
| 457 | VRC01 contact site | β23 (CD4 contacts) | Neutralization | 6 |
| 458 | VRC01 contact site | β23 (CD4 contacts) | Binding | 1 |
| 459 | VRC01 contact site | CD4 contact | Neutralization | 7 |
| 460 | VRC01 contact site | V5 Loop (CD4 contact) | Neutralization | 4 |
| 461 | VRC01 contact site | V5 Loop (CD4 contact) | Neutralization | 3 |
| 462 | VRC01 contact site | V5 Loop | Binding | 1 |
| 463 | VRC01 contact site | V5 Loop | Neutralization | 3 |
| 465 | VRC01 contact site | V5 Loop β24 | Binding | 1 |
| 469 | VRC01 contact site | V5 Loop β24 (CD4 contact) | Binding | 1 |
| 471 | Glycosylation site | β24 (CD4 contact) | Neutralization | 4 |
| 472 | VRC01 contact site | CD4 contact | Binding | 9 |
| 473 | VRC01 contact site | CD4 contact | Neutralization | 7 |
| 474 | VRC01 contact site | CD4 contact | Neutralization | 6 |

**References/Citations**


[1]Dingens, et al.2019. Immunity, Volume 50, Issue 2, 520 - 532.e3

[2]Juraska, et al., 2024, PNAS 121: e2308942121

[3]Falkowska, et al. J Virol. 2012;86(8):4394-4403

[4]LaBranche, et al. 2019 PLOS Pathog 14(11): e1007646

[5]Schommers, et al. Cell. 2020;180(3):471-489.e22

[6]Huang, et al. Immunity. 2016;45(5):1108-1121

[7]Cheng, et al. JCI Insight. 2018;3(5):e97018

[8]Foulkes, et al. PLoS Pathog. 2025;21(1):e1012825.

[9]Zhou, et al. Science. 2010;329(5993):811-817

**Table S2.** Metadata of participants with variant sites

| Trial | PtID | Treatment Arm | Parental (WT)/Mutation | VRC01 Sensitive/ Resistant | VRC01 infusion to TP1 (days) | TP1-TP2 (days) | No. Sequences Time point 1 | No. Sequences Time point 2 |
|---|---|---|---|---|---|---|---|---|
| 703 | 0520 | 10 mg/kg | WT | Sensitive | 0 | 28 | 84 | 291 |
| | | | D474N | Sensitive | | | 4 | 2 |
| 703 | 0578 | 10 mg/kg | WT | Sensitive | 0 | 9 | 137 | 153 |
| | | | N280D | Resistant | | | 25 | 131 |
| | | | G458E | Resistant | | | 7 | 22 |
| | | | N280D, G458E | Resistant | | | 0 | 1 |
| 703 | 0597 | 10 mg/kg | WT | Sensitive | 0 | 6 | 215 | 115 |
| | | | S461N | Sensitive | | | 89 | 62 |
| 703 | 0860 | 10 mg/kg | WT | Sensitive | 28 | 14 | 171 | 121 |
| | | | N276T | Resistant | | | 0 | 4 |
| | | | N276I | Resistant | | | 1 | 2 |
| 703 | 0967 | 10 mg/kg | WT | Sensitive | 81 | 8 | 42 | 23 |
| | | | N280D | Resistant | | | 0 | 5 |
| 703 | 1551 | 10 mg/kg | WT | Sensitive | 177 | 17 | 36 | 339 |
| | | | S365P | Sensitive | | | 0 | 13 |
| 703 | 1750 | Placebo | WT | Sensitive | No infusion | 8 | 252 | 361 |
| | | | V455I | Sensitive | | | 0 | 7 |
| 703 | 1783 | 10 mg/kg | WT | Sensitive | 0 | 9 | 361 | 393 |
| | | | N197T | Resistant | | | 0 | 17 |
| | | | D279A | Resistant | | | 0 | 4 |
| | | | K460N | Resistant | | | 0 | 26 |
| | | | N197T, K460N | Resistant | | | 0 | 1 |
| 703 | 2141 | 10 mg/kg | WT | Sensitive | 0 | 26 | 107 | 203 |
| | | | D279G | Resistant | | | 0 | 6 |
| 703 | 2372 | 10 mg/kg | WT | Sensitive | 0 | 23 | 374 | 192 |
| | | | D279G | Resistant | | | 0 | 91 |
| | | | D279Y | Resistant | | | 0 | 4 |
| | | | D279A | Resistant | | | 0 | 3 |
| | | | N280D | Resistant | | | 0 | 26 |
| | | | G459D | Resistant | | | 0 | 26 |
| | | | D279A, G459D | Resistant | | | 0 | 1 |
| | | | N280D, G459D | Resistant | | | 0 | 1 |
| 703 | 2631 | Placebo | WT | Sensitive | No infusion | 15 | 95 | 132 |
| | | | N461K | Sensitive | | | 2 | 5 |
| 704 | 0601 | 10 mg/kg | WT | Sensitive | 0 | 28 | 123 | 89 |
| | | | N279K | Resistant | | | 0 | 50 |
| 704 | 0855 | Placebo | WT | Sensitive | No infusion | 5 | 213 | 65 |
| | | | E370K | N/A* | | | 3 | 0 |
| 704 | 1481 | 10 mg/kg | WT | Sensitive | 27 | 8 | 379 | 357 |
| | | | D279Y | Resistant | | | 93 | 190 |
| 704 | 1551 | 10 mg/kg | WT | Sensitive | 0 | 6 | 182 | 30 |
| | | | P369T | Sensitive | | | 9 | 1 |
| | | | P369S | Sensitive | | | 1 | 0 |
| | | | T455I | Sensitive | | | 6 | 0 |
| | | | T455A | Sensitive | | | 1 | 1 |
| | | | R456K | Sensitive | | | 2 | 1 |
| | | | D462Y | Sensitive | | | 2 | 0 |
| | | | D462G | Sensitive | | | 1 | 0 |
| 704 | 1783 | Placebo | WT | Sensitive | No infusion | 20 | 395 | 264 |
| | | | D461N | Sensitive | | | 1 | 4 |
| 704 | 1802 | 10 mg/kg | WT | Sensitive | -27 | 27 | 297 | 90 |
| | | | D474N | Sensitive | | | 0 | 6 |
| 704 | 2063 | Placebo | WT | Sensitive | No infusion | 28 | 297 | 90 |
| | | | G459E | Sensitive | | | 0 | 7 |

*Pseudoviruses with E370K were not functional.

PtID, Participant study identifier, WT - wildtype

**Table S3: Correlation co-efficients calculated from random effects generalized linear model of Envelope backbones**

| bNAbs | | Corr. Coeff. (95% CI) | Pvalue |
|---|---|---|---|
| VRC01 | VRC01.23LS | 0.95 (0.74, 1) | 9.7E-11 |
| | VRC07-523LS | 0.84 (0.56, 1) | 2.3E-07 |
| | 3BNC117 | 1 (0.86, 1) | 1.9E-14 |
| | N6 | 0.32 (0.05, 0.59) | 0.02 |
| | 1-18 | -0.12 (-0.36, 0.14) | 0.36 |
| | CH235.12 | 0.13 (-0.19, 0.47) | 0.41 |
| | VRC_CH31 | 0.96 (0.79, 1) | 1.3E-15 |
| | PG9 | 0.09 (-0.17, 0.35) | 0.49 |
| | 10E8.v4 | -0.21 (-0.47, 0.05) | 0.11 |
| VRC01.23LS | VRC07-523LS | 0.66 (0.48, 0.85) | 1.9E-09 |
| | 3BNC117 | 0.49 (0.26, 0.76) | 8.5E-05 |
| | N6 | 0.29 (0.12, 0.48) | 1.8E-03 |
| | 1-18 | 0.1 (-0.06, 0.28) | 0.21 |
| | CH235.12 | 0.24 (0.02, 0.46) | 0.03 |
| | VRC_CH31 | 0.5 (0.31, 0.72) | 2.4E-06 |
| | PG9 | 0.05 (-0.12, 0.23) | 0.52 |
| | 10E8.v4 | -0.09 (-0.27, 0.09) | 0.32 |
| VRC07-523LS | 3BNC117 | 0.54 (0.3, 0.79) | 3.8E-05 |
| | N6 | 0.49 (0.33, 0.65) | 1.0E-07 |
| | 1-18 | 0.08 (-0.1, 0.28) | 0.37 |
| | CH235.12 | 0.24 (-0.01, 0.51) | 0.06 |
| | VRC_CH31 | 0.47 (0.27, 0.66) | 1.4E-05 |
| | PG9 | 0.04 (-0.14, 0.24) | 0.63 |
| | 10E8.v4 | 0.01 (-0.18, 0.22) | 0.87 |
| 3BNC117 | N6 | 0.12 (-0.1, 0.36) | 0.28 |
| | 1-18 | -0.18 (-0.36, 0.03) | 0.09 |
| | CH235.12 | 0.16 (-0.09, 0.42) | 0.21 |
| | VRC_CH31 | 0.76 (0.63, 0.88) | 2.1E-17 |
| | PG9 | 0.05 (-0.15, 0.25) | 0.62 |
| | 10E8.v4 | -0.16 (-0.36, 0.05) | 0.13 |
| N6 | 1-18 | 0.45 (0.23, 0.71) | 1.7E-04 |
| | CH235.12 | 0.49 (0.12, 0.97) | 0.01 |
| | VRC_CH31 | 0.28 (-0.02, 0.58) | 0.06 |
| | PG9 | 0.12 (-0.14, 0.39) | 0.36 |
| | 10E8.v4 | 0.22 (-0.04, 0.5) | 0.10 |
| 1-18 | CH235.12 | 0.65 (0.38, 0.92) | 3.9E-05 |
| | VRC_CH31 | 0.1 (-0.23, 0.42) | 0.46 |
| | PG9 | 0.2 (-0.09, 0.5) | 0.17 |
| | 10E8.v4 | 0.41 (0.15, 0.69) | 3.3E-03 |
| CH235.12 | VRC_CH31 | 0.2 (-0.05, 0.45) | 0.11 |
| | PG9 | 0.16 (-0.06, 0.4) | 0.16 |
| | 10E8.v4 | 0.27 (0.03, 0.52) | 0.03 |
| VRC_CH31 | PG9 | 0.07 (-0.15, 0.31) | 0.51 |
| | 10E8.v4 | -0.13 (-0.37, 0.11) | 0.29 |
| PG9 | 10E8.v4 | 0.55 (0.29, 0.78) | 3.1E-05 |
