## Supplementary Figures for "VRC01 Selects Rare HIV Escape Mutations After Acquisition in Antibody-Mediated Prevention Trials"

A) V703_0520 VRC01 10 mg/kg

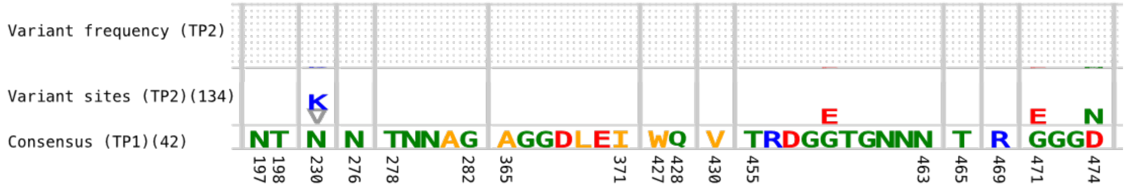


B) V703_0578 VRC01 10 mg/kg

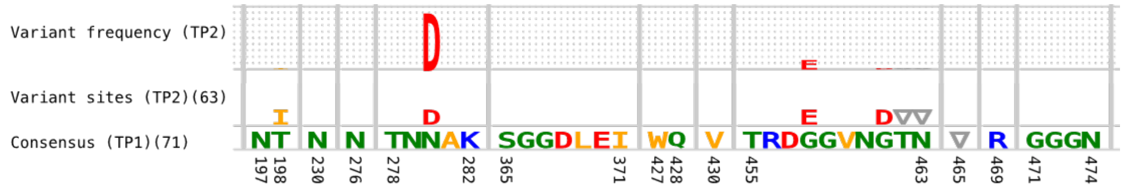


C) V703_0597 VRC01 10 mg/kg

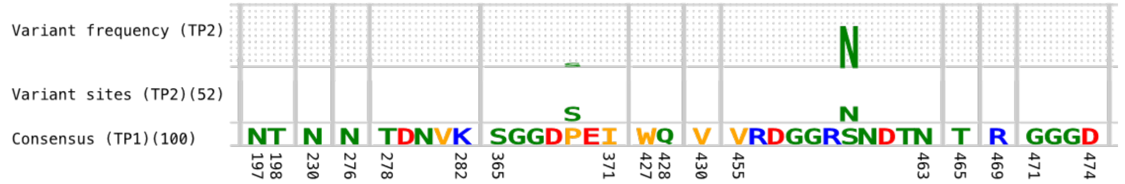


D) V703_0860 VRC01 10 mg/kg

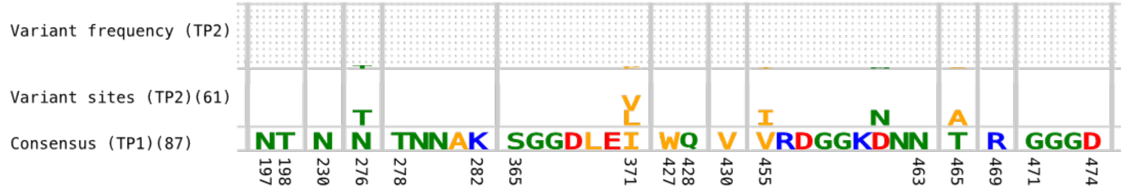


E) V703_0967 VRC01 10 mg/kg

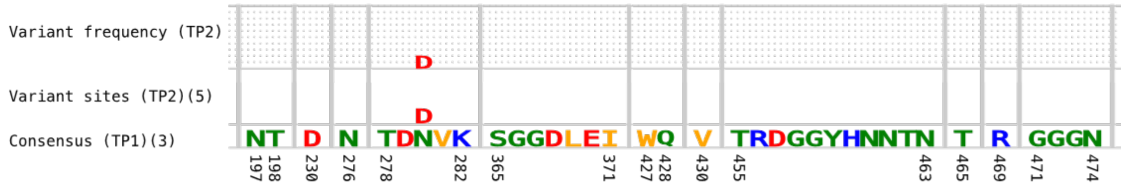


F) V703_1551 VRC01 10 mg/kg

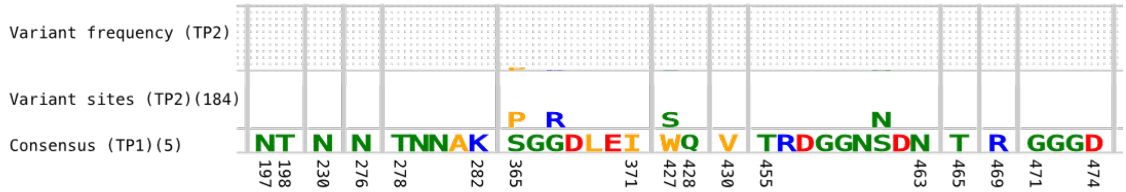


G) V703_1750 Placebo

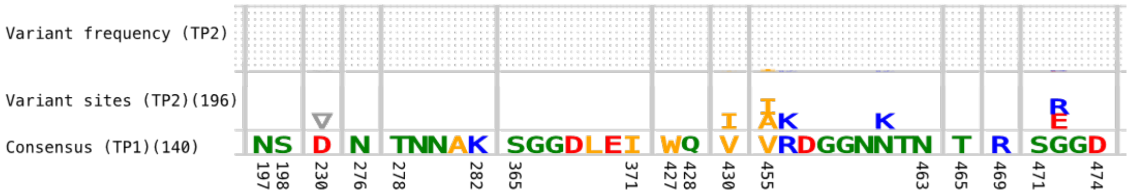

H) V703_1783 VRC01 10 mg/kg

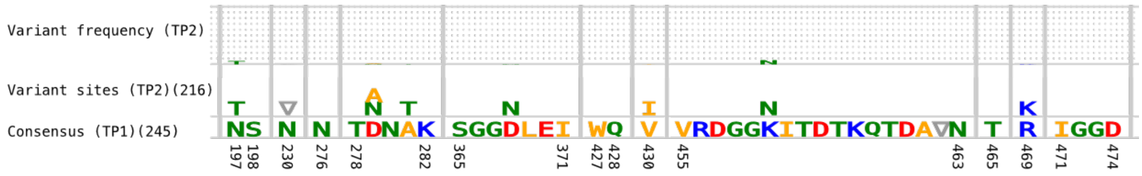


I) V703_2141 VRC01 10 mg/kg

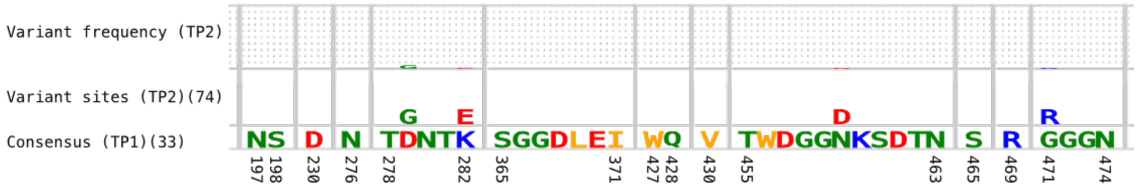


J) V703_2372 VRC01 10 mg/kg

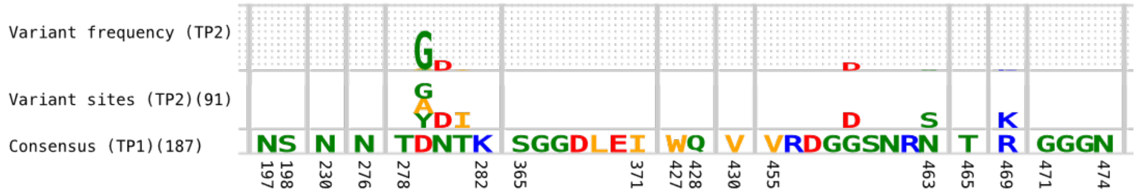


K) V703_2631 Placebo

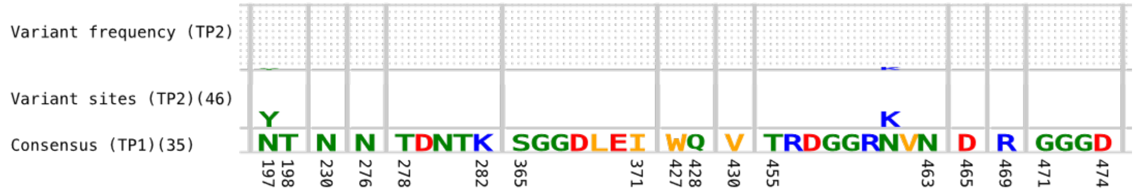


L) V704_0601 VRC01 10 mg/kg

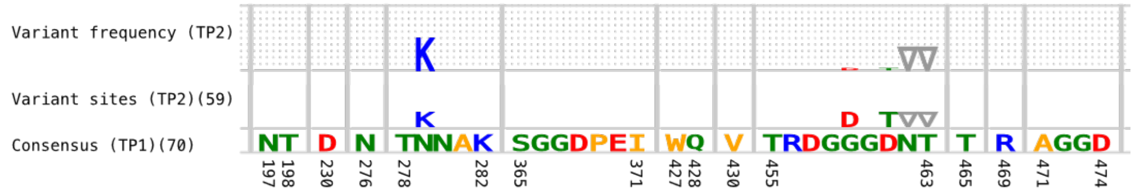


M) V704_0855 Placebo

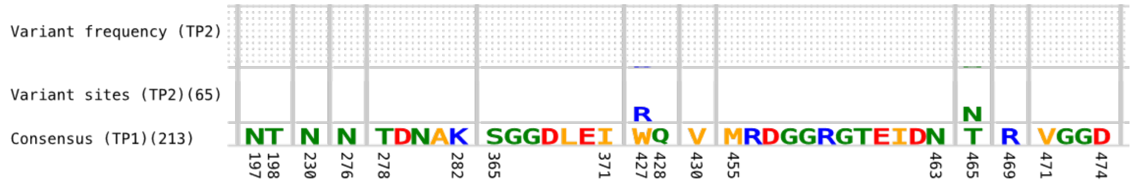


N) V704_1481 VRC01 10 mg/kg

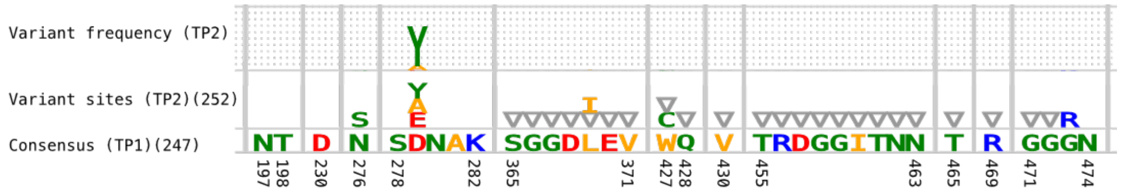

O) V704_1551 VRC01 10 mg/kg

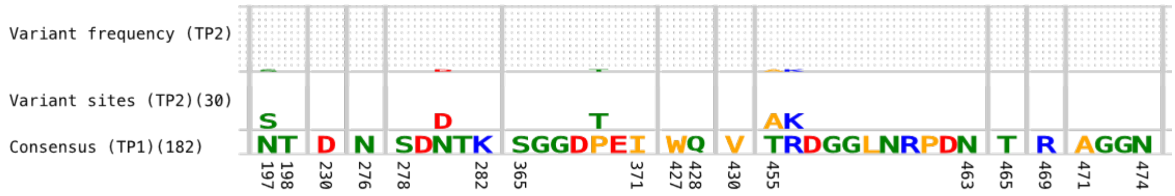


P) V704_1783 Placebo

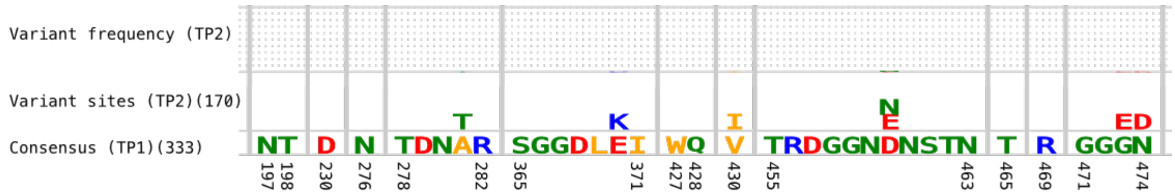


Q) V704_1802 VRC01 10 mg/kg

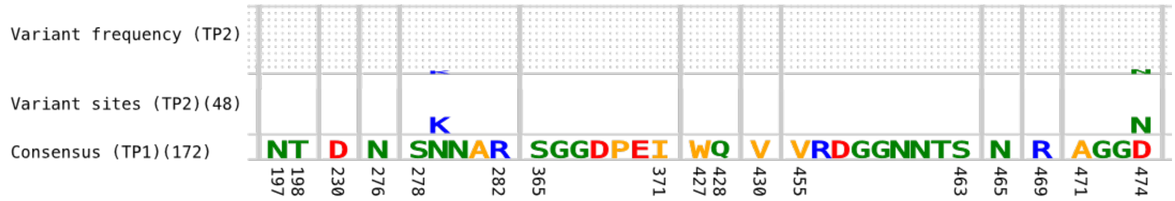


R) V704_2063 Placebo

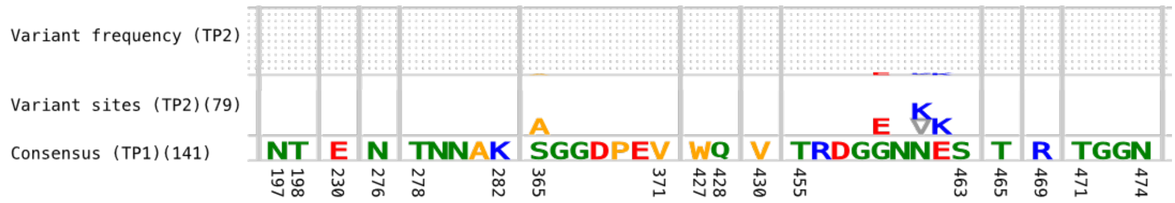


**Figure S1. Escape logograms for participants with variant sites in the 34 VRC01 epitope sites of interest.** HVTN 703/HPTN 081: A-K; HVTN704/HPTN 085 L-Q. Logograms showing amino acid (AA) changes between time point 1 (TP1) and time point 2 (TP2). The bottom panel presents the founder lineage consensus sequence at TP1, while the top panel shows the frequencies of variants at TP2. The middle panel highlights variant amino acids identified at a site compared to the consensus at TP1 and has been scaled for visibility. Sequence counts at each time point are shown in parentheses.

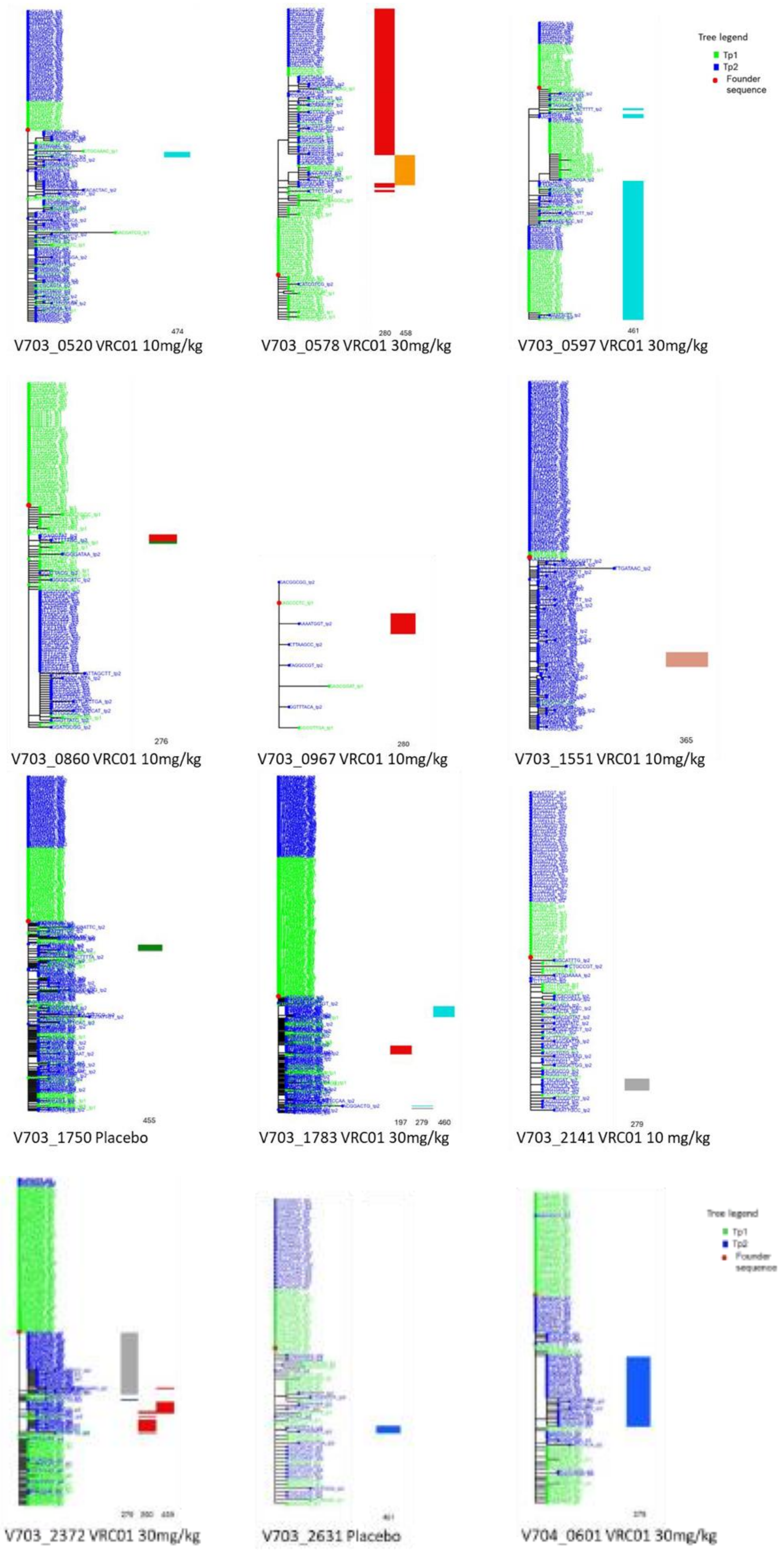

Tree legend
Tp1
Tp2
Founder sequence
474
V703_0520 VRC01 10mg/kg
280
458
V703_0578 VRC01 30mg/kg
461
V703_0597 VRC01 30mg/kg
276
V703_0860 VRC01 10mg/kg
280
V703_0967 VRC01 10mg/kg
365
V703_1551 VRC01 10mg/kg
455
V703_1750 Placebo
197
279
460
V703_1783 VRC01 30mg/kg
279
V703_2141 VRC01 10 mg/kg
V703_2372 VRC01 30mg/kg
V703_2631 Placebo
Tree legend
Tp1
Tp2
Founder sequence
V704_0601 VRC01 30mg/kg

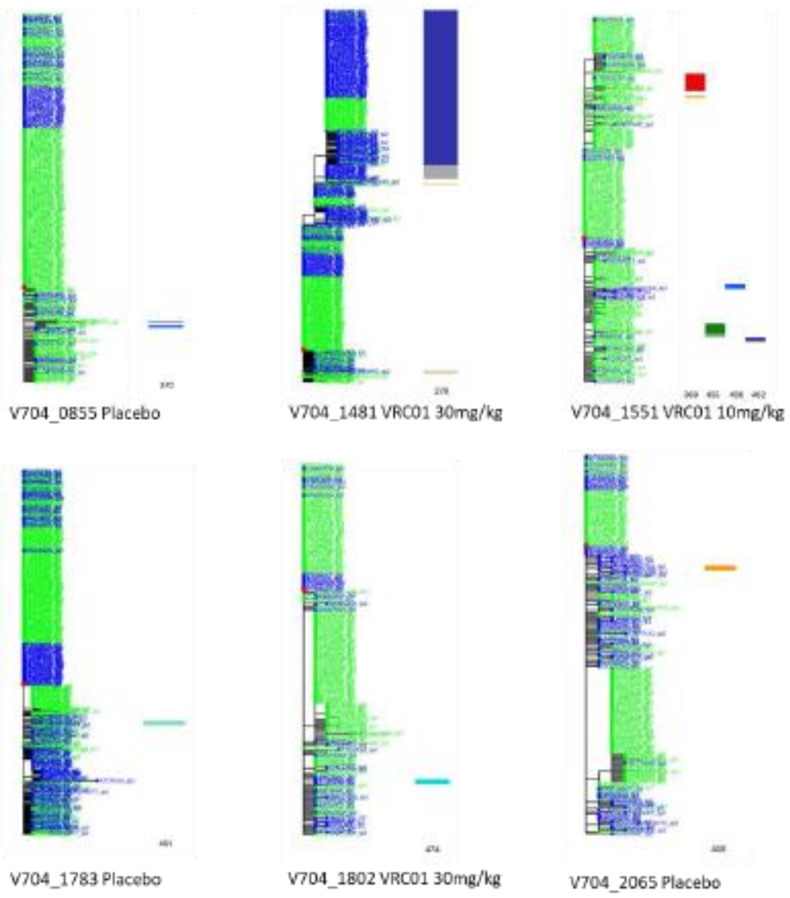


| Colour | AA name | Colour | AA name | Colour | AA name | Colour | AA name |
|---|---|---|---|---|---|---|---|
| | A – Alanine | | M – Methionine | | Y – Tyrosine | | P – Proline |
| | G - Glycine | | V – Valine | | S – Serine | | N – Asparagine |
| | I – Isoleucine | | F – Phenylalanine | | T – Threonine | | Q – Glutamine |
| | L – Leucine | | W – Tryptophan | | C – Cysteine | | K – Lysine |
| | R – Arginine | | H – Histidine | | D – Aspartate | | E - Glutamate |

**Figure S2.** Phylogenetic trees (left) displaying sequences from time point 1 (green) and time point 2 (blue). Adjacent highlighter plots show amino acid changes in the VRC01 epitope sites relative to a viral sequence (red dot) closest to the consensus at time point 1.

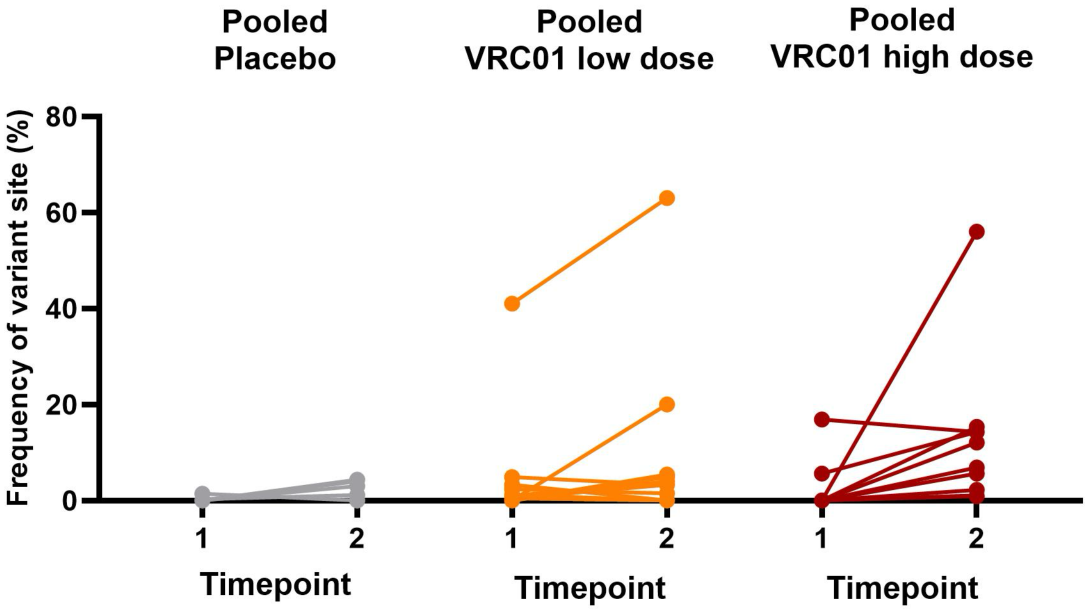


**Figure S3.** A graph indicating the change of prevalence of viral sequences with variant sites between sample time points. The median number of days between sampling time points: Placebo = 8 days (range 5-28), VRC01 low dose = 11 days (range 6-63) and VRC01 high dose =16 days (range 8-28).

| Treatment Arm | Virus | IC80 titer: CD4bs bnAbs: VRC01 | VRC01.23LS | VRC07-523LS | 3BNC117 | N6 | 1-18 | CH235.12 | VRC_CH31 | Controls: PG9 | 10E8.v4 |
|---|---|---|---|---|---|---|---|---|---|---|---|
| 10 mg/kg | V703_0520 WT | 0,56 | 0,06 | 0,21 | 0,2 | 0,19 | 0,31 | 25 | 0,57 | 6,46 | 12,1 |
| | D474N | 1,2 | 0,08 | 0,51 | 0,63 | 0,27 | 0,41 | 25 | 0,68 | 18,8 | 19,8 |
| 30 mg/kg | V703_0578 WT | 0,88 | 0,06 | 0,19 | 0,41 | 0,12 | 0,2 | 0,79 | 0,56 | 0,03 | 25 |
| | N280D | 25 | 25 | 25 | 25 | 0,31 | 0,1 | 25 | 25 | 0,01 | 3,54 |
| | G458E | 25 | 0,43 | 2,83 | 25 | 0,05 | 0,06 | 0,78 | 25 | 0,02 | 3,86 |
| | N280D G458E | 25 | 25 | 25 | 25 | 0,37 | 0,08 | 25 | 25 | 0,01 | 3,92 |
| 30 mg/kg | V704_0601 WT | 4,34 | 0,68 | 0,51 | 0,84 | 0,59 | 0,56 | 25 | 1,96 | 25 | 20,4 |
| | N279K | 25 | 0,13 | 25 | 25 | 6,15 | 0,03 | 0,56 | 25 | 25 | 23,3 |
| 10 mg/kg | V703_0860 WT | 5,14 | 0,89 | 0,35 | 0,28 | 0,5 | 0,15 | 7 | 1,92 | 25 | 4,9 |
| | N276T | 25 | 0,54 | 0,08 | 0,22 | 0,09 | 0,05 | 0,31 | 1,1 | 25 | 2,13 |
| | N276I | 25 | 25 | 0,25 | 6,03 | 0,14 | 0,03 | 0,27 | 25 | 25 | 0,52 |
| 10 mg/kg | V703_0967 WT | 0,13 | 0,28 | 0,03 | 0,07 | 0,03 | 0,06 | 0,06 | 0,33 | 0,02 | 0,29 |
| | N280D | 18 | 0,18 | 0,01 | 16,6 | 0,01 | 0,01 | 0,05 | 25 | 0,01 | 0,02 |
| 10 mg/kg | V703_1551 WT | 0,67 | 0,11 | 0,09 | 0,19 | 0,08 | 0,18 | 0,89 | 1,21 | 0,05 | 7,77 |
| | S365P | 0,23 | 0,04 | 0,03 | 0,04 | 0,02 | 0,07 | 0,09 | 0,56 | 0,03 | 10,2 |
| Placebo | V703_1750 WT | 3,2 | 0,7 | 0,71 | 1,05 | 0,79 | 2,04 | 8,64 | 21,6 | 25 | 25 |
| | V455I | 1,93 | 0,51 | 0,52 | 0,83 | 0,65 | 1,57 | 12,6 | 9,62 | 25 | 5,88 |
| 10 mg/kg | V704_1551 WT | 5,31 | 1 | 0,68 | 1,09 | 0,82 | 0,87 | 25 | 2,31 | 25 | 25 |
| | P369T | 2,81 | 0,17 | 0,6 | 0,76 | 1,41 | 1,2 | 25 | 1,4 | 1 | 3,09 |
| | P369S | 3,61 | 0,23 | 0,68 | 0,87 | 1,62 | 1,18 | 25 | 1,68 | 0,71 | 7,18 |
| | T455A | 1,81 | 0,14 | 0,35 | 0,95 | 0,85 | 1,08 | 25 | 1,29 | 0,68 | 4,84 |
| | T455I | 1,41 | 0,11 | 0,26 | 0,58 | 0,66 | 0,66 | 25 | 0,79 | 0,35 | 2,12 |
| | R456K | 0,89 | 0,09 | 0,19 | 0,41 | 0,5 | 0,65 | 12,3 | 0,61 | 0,41 | 2,18 |
| | D462G | 1,62 | 0,15 | 0,33 | 0,57 | 0,72 | 0,9 | 25 | 0,77 | 0,54 | 4,06 |
| | D462Y | 2,12 | 0,18 | 0,41 | 0,85 | 1,02 | 1,12 | 25 | 1,01 | 0,7 | 5,89 |
| 30 mg/kg | V703_1783 WT | 3,08 | 0,31 | 0,3 | 25 | 0,37 | 6,47 | 25 | 0,9 | 0,35 | 15,5 |
| | N197T K460N | 25 | 7,39 | 6,49 | 25 | 25 | 25 | 25 | 25 | 0,22 | 1,87 |
| | N197T | 25 | 1,89 | 1,87 | 25 | 2,57 | 25 | 25 | 25 | 0,09 | 4,89 |
| | D279A | 25 | 2,42 | 10 | 25 | 2,73 | 25 | 25 | 25 | 0,11 | 15,4 |
| | K460N | 19,8 | 2,48 | 1,85 | 25 | 2,02 | 25 | 25 | 25 | 0,76 | 21,6 |
| Placebo | V704_1783 WT | 0,85 | 0,15 | 0,55 | 0,42 | 0,59 | 0,09 | 25 | 1,94 | 25 | 25 |
| | D461N | 0,71 | 0,12 | 0,5 | 0,31 | 0,48 | 0,11 | 25 | 1,31 | 25 | 25 |
| 30 mg/kg | V704_1802 WT | 4,32 | 1,26 | 0,65 | 1,21 | 0,82 | 1,55 | 25 | 5,73 | 25 | 25 |
| | D474N | 2,42 | 0,18 | 0,5 | 0,73 | 1,38 | 0,96 | 25 | 1,76 | 25 | 19 |
| Placebo | V704_2065 WT | 0,41 | 0,07 | 0,18 | 0,15 | 0,1 | 0,16 | 0,66 | 1,78 | 0,38 | 0,55 |
| | G459E | 1,04 | 0,04 | 0,16 | 2,24 | 0,09 | 0,08 | 0,7 | ND | 0,53 | 0,54 |
| 10 mg/kg | V703_2141 WT | 5,04 | 0,32 | 0,82 | 17,9 | 0,74 | 1,2 | 25 | 25 | 4,9 | 25 |
| | D279G | 25 | 25 | 25 | 25 | 25 | 25 | 11,5 | 25 | 1,74 | 5,93 |
| 30 mg/kg | V703_2372 WT | 2,74 | 0,3 | 0,56 | 0,8 | 1,13 | 0,21 | 6,29 | 2,35 | 25 | 0,15 |
| | N280D | 25 | 0,59 | 17,7 | 25 | 0,27 | 0,17 | 25 | 25 | 25 | 0,08 |
| | G459D | 25 | 0,33 | 0,7 | 25 | 0,32 | 0,14 | 4,34 | 25 | 25 | 0,08 |
| | D279G | 25 | 1,55 | 25 | 25 | 0,68 | 0,13 | 1,28 | 25 | 25 | 0,19 |
| | D279Y | 25 | 3,38 | 25 | 25 | 0,21 | 0,06 | 0,7 | 25 | 25 | 0,07 |
| | D279A | 25 | 1 | 25 | 1,21 | 0,87 | 0,17 | 1,81 | 6,43 | 25 | 0,08 |
| | D279A G459D | 25 | 25 | 25 | 25 | 0,48 | 0,17 | 1,7 | 25 | 25 | 0,12 |
| | N280D G459D | 25 | 25 | 25 | 25 | 0,12 | 0,05 | 25 | 25 | 25 | 0,04 |
| Placebo | V703_2631 WT | 3,02 | 0,6 | 0,65 | 1,12 | 1,41 | 0,64 | 11,4 | 10,7 | 0,29 | 25 |
| | N461K | 0,65 | 0,13 | 0,14 | 0,14 | 0,24 | 0,11 | 0,44 | 0,81 | 0,01 | 0,42 |

| Treatment Arm | Virus | Fold difference (Mutant/WT): CD4bs bnAbs: VRC01 | VRC01.23LS | VRC07-523LS | 3BNC117 | N6 | 1-18 | CH235.12 | VRC_CH31 | Controls: PG9 | 10E8.v4 |
|---|---|---|---|---|---|---|---|---|---|---|---|
| 10 mg/kg | V703_0520 WT | | | | | | | | | | |
| | D474N | 0,35 | 1,3 | 2,42 | 3,15 | 1,42 | 1,32 | 1 | 1,19 | 2,9 | 1,63 |
| 30 mg/kg | V703_0578 WT | | | | | | | | | | |
| | N280D | 28,4 | 417 | 132 | 61 | 2,58 | 0,5 | 31,6 | 44,6 | 0,33 | 0,14 |
| | G458E | 28,4 | 7,17 | 14,9 | 61 | 0,42 | 0,3 | 0,99 | 44,6 | 0,67 | 0,15 |
| | N280D G458E | 28,4 | 417 | 132 | 61 | 3,08 | 0,4 | 31,6 | 44,6 | 0,33 | 0,16 |
| 30 mg/kg | V704_0601 WT | | | | | | | | | | |
| | N279K | 5,76 | 0,19 | 49 | 29,8 | 10,4 | 0,05 | 0,02 | 12,8 | 1 | 1,14 |
| 10 mg/kg | V703_0860 WT | | | | | | | | | | |
| | N276T | 4,86 | 0,61 | 0,23 | 0,79 | 0,18 | 0,33 | 0,04 | 0,57 | 1 | 0,43 |
| | N276I | 4,86 | 28,1 | 0,71 | 21,5 | 0,28 | 0,2 | 0,04 | 13 | 1 | 0,11 |
| 10 mg/kg | V703_0967 WT | | | | | | | | | | |
| | N280D | 138 | 0,64 | 0,17 | 238 | 0,17 | 0,08 | 0,83 | 75,8 | 0,5 | 0,07 |
| 10 mg/kg | V703_1551 WT | | | | | | | | | | |
| | S365P | 0,34 | 0,36 | 0,33 | 0,21 | 0,25 | 0,39 | 0,1 | 0,46 | 0,6 | 1,31 |
| Placebo | V703_1750 WT | | | | | | | | | | |
| | V455I | 0,6 | 0,73 | 0,73 | 0,79 | 0,82 | 0,77 | 1,46 | 0,45 | 1 | 0,2 |
| 10 mg/kg | V704_1551 WT | | | | | | | | | | |
| | P369T | 0,53 | 0,17 | 0,88 | 0,7 | 1,72 | 1,38 | 1 | 0,61 | 0,04 | 0,17 |
| | P369S | 0,68 | 0,23 | 1 | 0,8 | 1,98 | 1,36 | 1 | 0,73 | 0,03 | 0,29 |
| | T455A | 0,34 | 0,14 | 0,51 | 0,87 | 1,04 | 1,24 | 1 | 0,56 | 0,03 | 0,19 |
| | T455I | 0,27 | 0,11 | 0,38 | 0,53 | 0,8 | 0,76 | 1 | 0,34 | 0,01 | 0,08 |
| | R456K | 0,17 | 0,09 | 0,28 | 0,38 | 0,61 | 0,75 | 0,49 | 0,26 | 0,02 | 0,09 |
| | D462G | 0,31 | 0,15 | 0,49 | 0,52 | 0,88 | 1,03 | 1 | 0,33 | 0,02 | 0,16 |
| | D462Y | 0,4 | 0,18 | 0,6 | 0,78 | 1,24 | 1,29 | 1 | 0,44 | 0,03 | 0,24 |
| 30 mg/kg | V703_1783 WT | | | | | | | | | | |
| | N197T K460N | 8,12 | 23,8 | 21,6 | 1 | 67,6 | 3,87 | 1 | 27,8 | 0,63 | 0,12 |
| | N197T | 8,12 | 6,1 | 6,23 | 1 | 6,95 | 3,87 | 1 | 27,8 | 0,26 | 0,32 |
| | D279A | 8,12 | 7,81 | 33,3 | 1 | 7,38 | 3,87 | 1 | 27,8 | 0,31 | 0,99 |
| | K460N | 6,42 | 8 | 6,17 | 1 | 5,46 | 3,87 | 1 | 27,8 | 2,17 | 1,39 |
| Placebo | V704_1783 WT | | | | | | | | | | |
| | D461N | 0,84 | 0,8 | 0,91 | 0,74 | 0,81 | 1,22 | 1 | 0,68 | 1 | 1 |
| 30 mg/kg | V704_1802 WT | | | | | | | | | | |
| | D474N | 0,56 | 0,14 | 0,77 | 0,6 | 1,68 | 0,62 | 1 | 0,31 | 1 | 1,3 |
| Placebo | V704_2065 WT | | | | | | | | | | |
| | G459E | 2,54 | 0,5 | 0,89 | 14,9 | 0,9 | 0,5 | 1,05 | | 1,38 | 0,97 |
| 10 mg/kg | V703_2141 WT | | | | | | | | | | |
| | D279G | 4,96 | 78,1 | 30,5 | 1,4 | 33,8 | 20,8 | 0,46 | 1 | 0,36 | 0,24 |
| 30 mg/kg | V703_2372 WT | | | | | | | | | | |
| | N280D | 9,12 | 1,97 | 31,6 | 31,3 | 0,24 | 0,81 | 3,97 | 10,6 | 1 | 0,53 |
| | G459D | 9,12 | 1,1 | 1,25 | 31,3 | 0,28 | 0,67 | 0,69 | 10,6 | 1 | 0,53 |
| | D279G | 9,12 | 5,17 | 44,6 | 31,3 | 0,6 | 0,62 | 0,2 | 10,6 | 1 | 1,27 |
| | D279Y | 9,12 | 11,3 | 44,6 | 31,3 | 0,19 | 0,29 | 0,11 | 10,6 | 1 | 0,47 |
| | D279A | 9,12 | 3,33 | 44,6 | 1,51 | 0,77 | 0,81 | 0,29 | 2,74 | 1 | 0,53 |
| | D279A G459D | 9,12 | 83,3 | 44,6 | 31,3 | 0,42 | 0,81 | 0,27 | 10,6 | 1 | 0,8 |
| | N280D G459D | 9,12 | 83,3 | 44,6 | 31,3 | 0,11 | 0,24 | 3,97 | 10,6 | 1 | 0,27 |
| Placebo | V703_2631 WT | | | | | | | | | | |
| | N461K | 0,22 | 0,22 | 0,22 | 0,13 | 0,17 | 0,17 | 0,04 | 0,08 | 0,03 | 0,02 |

Keys

IC80 titres
- <0.1
- >0.1-1
- >1-10
- >10-25

Fold change
- >300
- >100-300
- >21-100
- >3-20
- <3

**Figure S4.** Heat plot of $IC_{80}$ neutralization titers of pseudoviruses and corresponding mutants. The plot on the left illustrating the $IC_{80}$ titers, and the grayscale plot on the right showing the fold difference between the sensitive PSV and the mutant. The eight CD4bs bnAbs are shown on the top, together with two control bnAbs targeting the V2 apex (PG9) and MPER (10E8v4) regions.

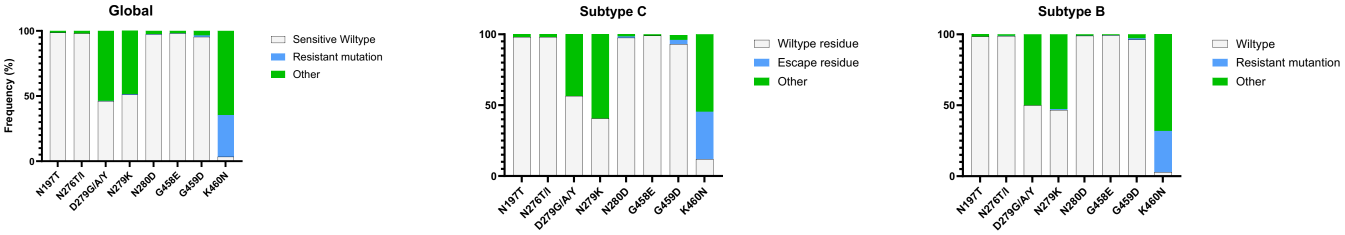


**Figure S5.** Frequencies of wildtype residues (AA in the transmitted founder lineage, light grey), escape residues (blue), other (neither wildtype nor escape, dark grey). **A.** HIV Group M sequences (n=8,452); **B**. Subtype C sequences (n=1779); **C.** Subtype B sequences (n=3043). Filtered sequence alignments downloaded from the Los Alamos database (https://www.hiv.lanl.gov/)
